# The legacy of past climate warming: strong local adaptation in rear-edge populations

**DOI:** 10.1101/2025.08.15.669932

**Authors:** Antoine Perrier, Olivia J. Keenan, Jeremiah W. Busch, Laura F. Galloway

## Abstract

Improving forecasts of species’ responses to climate change has become a central challenge in ecology and evolution as species distributions are increasingly disrupted by ongoing climate warming. Insight into this challenge may be gained through a better understanding of evolutionary responses to past climate change. The rear edges of species’ distributions are typically relict populations persisting in former glacial refugia at warmer range limits. As such, they form natural laboratories to study the evolutionary outcomes of past climate warming. Three such outcomes have been proposed for rear-edge populations: maintenance of high diversity due to long-term persistence, strong genetic drift following habitat decline, and strong local adaptation allowing persistence despite environmental change. Empirical studies rarely explicitly test these alternate outcomes limiting our understanding of evolutionary responses to warming climates. We tested all three evolutionary outcomes at the rear edge of the North American herb *Campanula americana*, by assessing genetic variation in a genome-wide population genetic study, drift load as measure of fitness decline due to drift in a controlled crossing study, and local adaptation in a transplant study. Rear-edge populations exhibited reduced genetic diversity within populations and high differentiation among populations, typically interpreted as evidence of genetic drift, yet show limited drift load. Instead, these populations expressed strong local adaptation, thriving in rear-edge habitats that were unsuitably warm for populations in the expanded range. This indicates that warm-edge populations may persist under warming climates by gradually adapting, even in the face of genetic erosion. Our findings highlight the importance of explicitly testing for all evolutionary outcomes, and particularly going beyond measures of genetic variation, when inferring evolutionary history. More broadly, these findings identify rear-edge populations not as relics of decline, but as underappreciated models for studying successful adaptation under long-term climate change.

## Introduction

Anticipating global shifts in biodiversity under contemporary climate change (Lenoir & Svenning, 2015; Rubenstein et al., 2023; Urban, 2024) is a central challenge in ecology and evolution. Ecological models generally predict shifts in distribution toward higher latitudes and elevations as organisms track suitable habitats (Parmesan & Yohe, 2003), with high risks of extinctions at warmer range limits (Cahill et al., 2014). While these predictions are broadly supported in empirical studies of the responses to ongoing climate change (e.g. Anderson et al., 2025), many exceptions have been documented (Alexander et al., 2018; Duchenne et al., 2021; Geppert et al., 2020; Rubenstein et al., 2020, 2023; Vilà-Cabrera et al., 2019). These exceptions highlight the need for a better fundamental understanding of how species respond to changing environments. Integrating evolutionary processes into projections of environmental change (Aguirre-Liguori et al., 2021; Nadeau & Urban, 2019) may improve predictions of species’ responses to climate change. Insights into these processes may be gained by studying how past climate change, particularly warming since the Last Glacial Maximum (LGM, ∼20kya; Hughes et al., 2013), has shaped contemporary ecological and evolutionary patterns across populations (Davis & Shaw, 2001; de Lafontaine et al., 2018; G. Hewitt, 2000; G. M. Hewitt, 2004).

Rear-edge populations, typically glacial relicts persisting at warmer range limits, can serve as natural laboratories to study evolutionary processes associated with climate change (Hampe & Petit 2005; Nadeau & Urban 2019; Perrier et al., *in review*). During the LGM, many temperate species were constrained to glacial refugia at low latitudes and elevations. As the Earth warmed, they expanded their ranges by tracking shifts in suitable habitats (“leading edges”; Hewitt 2000, 2004). In some species, refugial populations persisted more-or-less in place despite substantial warming, and now form the “rear edge” of the species distribution (“stable edge” sensu Hampe & Petit 2005). The environmental history of these populations offers a unique opportunity to study how warming climates may alter patterns of genetic diversity and structure, affect population demography, and impose new selective pressures. A better understanding of these past evolutionary responses will also be key in refining predictions of vulnerability and evolutionary potential under future warming at the rear edge (Perrier et al., *in review*).

Evolutionary responses to past warming can be inferred from genetic patterns in contemporary populations. Theoretical and empirical work on rear-edge populations identify three main genetic patterns, each representing a distinct outcome of evolution under past climate change (reviewed in Perrier et al., *in review*). The first predicted outcome is that rear edges are hotspots of genetic variation within and among populations. Rear edges may retain ancestral variation and thus have greater genetic diversity than populations established during postglacial colonization that often involve serial bottlenecks and founder effects (Excoffier et al., 2009). In addition, the older age and potential long-term isolation in shrinking suitable habitat may contribute to higher differentiation among rear-edge populations than younger populations in the expanded range (Hampe & Petit, 2005). A recent review supports this, finding substantially higher differentiation among rear-edge populations than in the expanded range (Perrier et al., *in review*). However, only about half of surveyed studies reported the expected greater genetic diversity within rear-edge populations, indicating that simple expectations fail to explain patterns of genetic variation.

The second predicted evolutionary outcome of past climate warming is the potential for strong genetic drift in rear-edge populations. Postglacial warming is expected to have caused substantial habitat loss and fragmentation at warmer range limits (Vilà-Cabrera et al., 2019), resulting in long-term demographic decline and isolation of populations. These processes may yield strong genetic drift in rear-edge populations, eroding diversity (Frankham, 1996; Young et al., 1996), reducing adaptive potential (Leonardi et al., 2012; Willi et al., 2006) and causing fitness decline due to the accumulation of deleterious mutations (i.e. drift load; Kimura et al., 1963; Nei et al., 1975; Lynch et al., 1995a, b; Kirkpatrick & Jarne 2000). Rear-edge populations often have low within-population diversity but high between population differentiation (e.g. Scalfi et al., 2009; Diekmann & Serrão 2012; Assis et al., 2014; Carbognani et al., 2019; Kebaïli et al., 2022), commonly interpreted as a signature of genetic drift (Eckert et al., 2008; Pironon et al., 2017; Perrier et al., *in review*). However, this pattern provides only indirect evidence of drift, and does not address whether drift affects fitness and adaptive potential. We lack studies that explicitly measure drift, e.g. using drift load (but see Perrier et al., 2020; Willi et al., 2018), that would yield conclusive evidence that environmental decline and demographic contraction under past warming has resulted in genetic drift in rear-edge populations.

The third predicted evolutionary outcome of past climate warming at the rear edge is strong local adaptation resulting from long-term directional selection in post-glacial climates (Bontrager et al., 2021). This outcome suggests that populations persist not as remnants, destined for decline, but because they have adapted and can exploit the warmer conditions. The few studies that have tested this possibility found heightened genomic signatures of selection (Keller et al., 2018; Parisod & Joost, 2010) and local adaptation (Mathiasen & Premoli, 2016; Saada et al., 2016) in rear-edge populations. Furthermore, a recent meta-analysis found that local adaptation increases toward equatorial range limits, often home to rear-edge populations (Bontrager et al., 2021), implying that strong local adaptation in rear edges may be more common than currently appreciated. Finally, rear-edge populations often show specializations to warm environments suggesting local adaptation (e.g. Ghouil et al., 2020; Pelletier et al., 2023; Perrier et al., 2025a). While many lines of evidence support the persistence of rear-edge populations under climate warming through adaptation, limited direct tests leave this possibility relatively underexplored and potentially underestimated.

Understanding how past climate change has shaped contemporary ecological and evolutionary patterns in rear edges requires evaluating all three outcomes – diversity, drift, and local adaptation. We test these outcomes by integrating climate, genetic, and experimental data on rear-edge populations of the herb *Campanula americana.* We first evaluate whether rear-edge populations are hotspots of genetic variation by assessing patterns of genetic diversity within and differentiation between populations (Study 1). We then test for genetic drift by estimating the expression of drift load in a crossing experiment (Study 2). Finally, we estimate local adaptation in a large-scale common garden experiment (Study 3). All studies include populations spanning the range to enable direct comparisons between rear-edge populations and populations that resulted from postglacial range expansion (hereafter “expanded range”). We hypothesize that rear-edge populations will have greater genetic diversity within and among populations, express greater drift load and/or demonstrate more local adaptation than populations from expanded regions. These complementary approaches provide a holistic framework using rear-edge populations as models to decipher how populations evolve when climates warm. This research also bridges critical gaps in our understanding of the consequences of past warming for contemporary populations, providing key insights for anticipating their responses to future change.

## Material and Methods

### Study system

*Campanula americana* is a monocarpic herb found in open forested areas across the eastern USA (Fig. 1A). Seeds typically germinate in spring or fall and survive over winter as rosettes (Baskin & Baskin, 1984; Galloway, 2002). Exposure to cold (*i.e.* vernalization, Chouard 1960) is essential to cue reproduction (Perrier et al., 2025a), that is initiated in spring by the elongation of the rosettes into a stalk, i.e. bolting. Flower buds develop and start to open in mid-summer, with the bulk of flowers opening progressively over ca. five weeks. Fruit maturation takes ca. one month and occurs from late summer to early fall.

**Figure 1:**
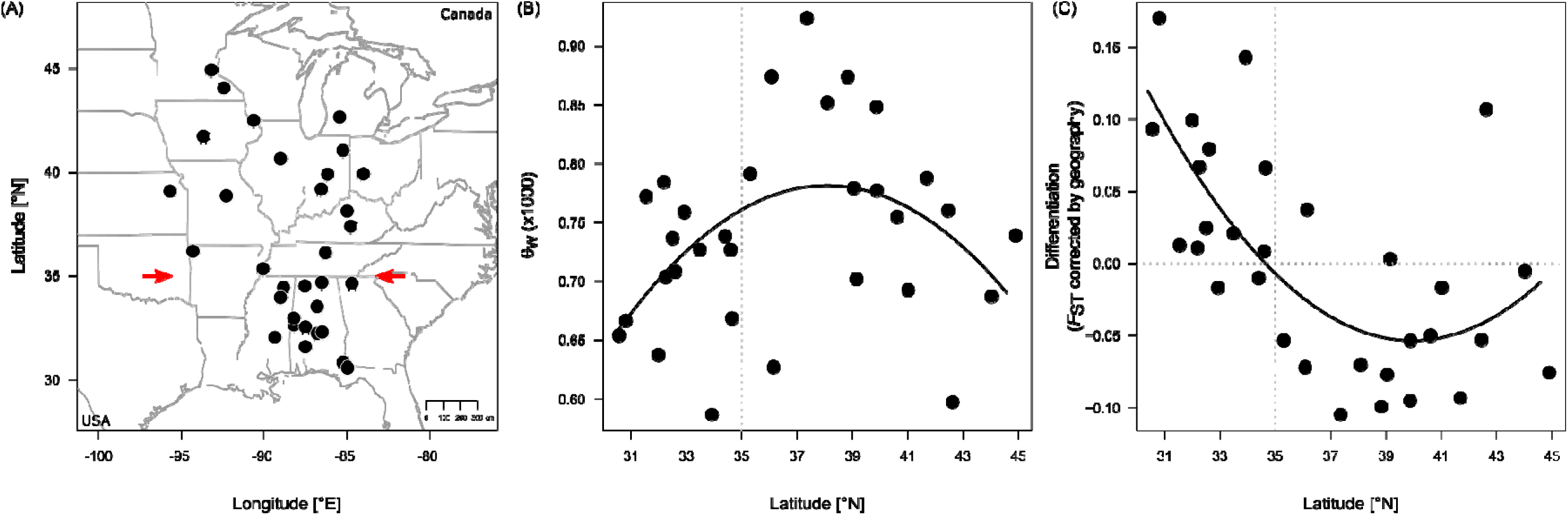
Variation in diversity and differentiation across the range (Study 1) **(A)** Location of the sequenced populations (dots) used to estimate genetic diversity and differentiation, with the range of the Western lineage indicated (light gray). Red arrows represent the latitudinal delimitation of the rear edge. **(B**) Genetic diversity within populations based on Watterson’s θ_W_ and **(C)** population-average differentiation among populations (*F*_ST_ after accounting for geographic distance) estimated for each population (dots). In (C), the horizontal dotted line indicates whether differentiation is higher (>0) or lower (<0) than expected based on isolation by distance. Vertical dotted lines represent the separation between the rear edge (<35°N) and the expanded range. Black lines represent the significant model-predicted relationship between each estimate and population latitude, with the 95% confidence interval indicated by shading. Test statistics are reported in Table 1.

**Table 1:** Test of variation in genetic diversity (D_π_ **and** D_W_) and differentiation (Study 1), and drift load (Study 2), across the range of *Campanula americana*.

| Dependent variable | N | Latitude |  | F |  | R<sup>2</sup> |
| --- | --- | --- | --- | --- | --- | --- |
| | | $\beta_{\text{linear}}$ | $\beta_{\text{square}}$ | | | |
| $\square_w$ | 31 | **8.05e-05** | **-1.82e-04** | **3.46** | * | 0.14 |
| $\square_{\pi}$ | 31 | -2.98e-05 | -8.93e-05 | 1.97 | | 0.06 |
| Differentiation | 31 | **-2.35e-01** | **1.59e-01** | **12.65** | *** | 0.44 |
| Drift load | 20 | **1.65e-02** | - | **22.37** | *** | 0.54 |

*Campanula americana* occupies a large geographic and climatic range, spanning a ∼ 1700km latitudinal gradient from subtropical areas close to the Gulf coast to the Great Lakes, and a ∼1800km longitudinal gradient from just west of the Mississippi to the Appalachian Mountains. Populations across the range belong to three geographically distinct genetic clades that diverged ca. 0.7 – 7.0 mya (Barnard-Kubow et al., 2015), and come into contact along the Appalachian mountains (Lamb et al., 2024). Here we focus on the “Western” clade, which includes most of the species’ populations (∼ 80%), and is found at low elevation (<600m) west of the Appalachian Mountains (Fig. 1A). Known hybrid zones between lineages (Lamb et al., 2024) were excluded from this study.

In this clade, the rear edge is geographically defined as the lower latitudinal third of the range (below 35° N, Fig. 1A, Perrier et al., 2025a). Here, populations overlap with putative glacial refugia and are ancestral in the clade (Barnard-Kubow et al., 2015), indicating a stable rear edge. Populations north of the rear edge are the result of postglacial range expansion from a mid-latitude origin estimated in eastern Kentucky (Koski et al., 2019; Prior et al., 2020). Contemporary rear-edge habitats are of lower suitability than the expanded range (Barnard-Kubow et al., 2015) and are substantially warmer, especially in winter (Perrier et al., 2025a), with reduced vernalization requirements and a distinctive delay in flowering (Perrier et al., 2025a).

### Study 1: Are rear edges reservoirs of diversity?

We assessed the hypothesis that rear-edge populations are hotspots of genetic variation by testing for differences in genetic diversity within populations and in genetic differentiation among populations across the species’ range.

#### Sequencing and genotyping

We sequenced 31 populations spanning the distribution of *C. americana*’s Western clade (Fig. 1A, Table S1). We collected leaf tissue from seven to ten individuals per population (292 total, Method S1, BioProject PRJNA1306192). DNA extraction was performed by the Genomics & Cell Characterization Core Facility at the University of Oregon (Eugene, OR, USA). Sequencing was performed using a Restriction-site Associated DNA sequencing (RAD-Seq) approach (Andrews et al., 2016; Baird et al., 2008). Library preparation and sequencing were performed by Floragenex Inc. (Beaverton, OR). More details of the plant material, DNA extraction, library preparation and sequencing are provided in Method S1.

Raw reads were processed into single nuclear polymorphism (SNP) calls using a modified STACKS v2 pipeline (Rochette et al., 2019), detailed in Method S1. In short, reads were demultiplexed, filtered and aligned to a haploid reference genome (A. Lopez-Caamal, *unpublished data*). Genotypes were called in diploid variant call format (VCF), then filtered for missingness (85%), minor allele count (2), sequencing depth (20x – 70x) and indels, for a final set of 137,312 SNPs. As this plant is an old tetraploid, diploid genotypes were converted into tetraploid genotypes using a custom R script (Method S1) adapted from a previous study in the species (Prior et al., 2020; code and VCF available in Perrier et al., 2025b).

#### Estimates of genetic variation

We estimated population-level genetic diversity as Watterson’s (θ_W_) and Tajima’s (θ_π_) estimators of θ (Tajima, 1989; Watterson, 1975). Both are measures of diversity, but θ_W_is more sensitive to rare variants (segregating sites) than θ_π_, which measures average pairwise differences. These estimates were calculated for each population in a sliding window approach using a custom R script (adapted from Koski et al., 2019, available in Perrier et al., 2025b), where each window was a distinct RAD locus (specific locations in the genome where RAD tags are derived from and shared across samples, Method S1), then averaged at the level of the population. We estimated differentiation among populations by first calculating pairwise F_ST_ (Weir & Cockerham, 1984) using the *StAMPP* package in R (Pembleton et al., 2013). To account for isolation-by-distance (IBD; Eckert et al., 2008; Slatkin 1993), we tested the relationship between linearized pairwise F_ST_ estimates (Rousset, 1997) and pairwise geographic distances, and used the residuals from this relationship to quantify population-level genetic differentiation relative to IBD expectations (hereafter “differentiation”, Methods S1).

To evaluate the expectation that rear edges are hotspots of genetic diversity, we assessed variation in population-level estimates of θ_W_, θ_π_, and differentiation in separate linear models that tested for the effect of a population’s latitude. Preliminary analysis evaluated whether latitude was best described by a linear or a second-degree quadratic relationship (c.f. Pironon et al., 2017), and results are reported for the best model based on AICc scores (Sugiura, 1978). If both models performed equally well (|ΔAICc| ≤ 2), the model with highest R^2^ was chosen. Dependent variables were tested assuming a normal distribution. For each model (and in all subsequent analyses), we tested and confirmed model assumptions.

### Study 2: Do rear-edge populations suffer from strong genetic drift?

We tested the hypothesis that rear-edge populations show elevated drift load, consistent with a history of strong genetic drift, by quantifying heterosis across the range in a crossing experiment. Heterosis, the increased fitness of between-population crosses (BPC) relative to within-population crosses (WPC), reflects the masking of recessive deleterious mutations presumably accumulated under drift in the parental populations (Crow, 1987).

#### Crossing design, raising of offspring and recording of traits

The experimental design was similar to a previous study investigating the accumulation of drift load in the expanded range of the species (Koski et al., 2019), to allow for comparison. In the present study, we focused on the rear edge and selected 19 populations spanning the rear edge and mid-latitudes to experimentally measure heterosis (Fig. 2, Table S2A). We extended the sampling to the leading edge by incorporating heterosis data from eight high-latitude populations (Fig. 2, Table S2A) generated in Koski et al., (2019). For each of the 19 rear-edge and mid-latitude populations, we raised one seedling from up to 18 independent seed families (mean 14.5 families) collected in nature in 2020 and 2021 to perform WPC and BPC crosses. Seedlings were raised under controlled conditions simulating the species’ seasonal cycle in 2021-2022 (Method S2). Germination and initial growth occurred in growth chambers simulating fall, followed by vernalization in a cold room simulating winter, and then reproduction in a greenhouse simulating summer.

**Figure 2:**
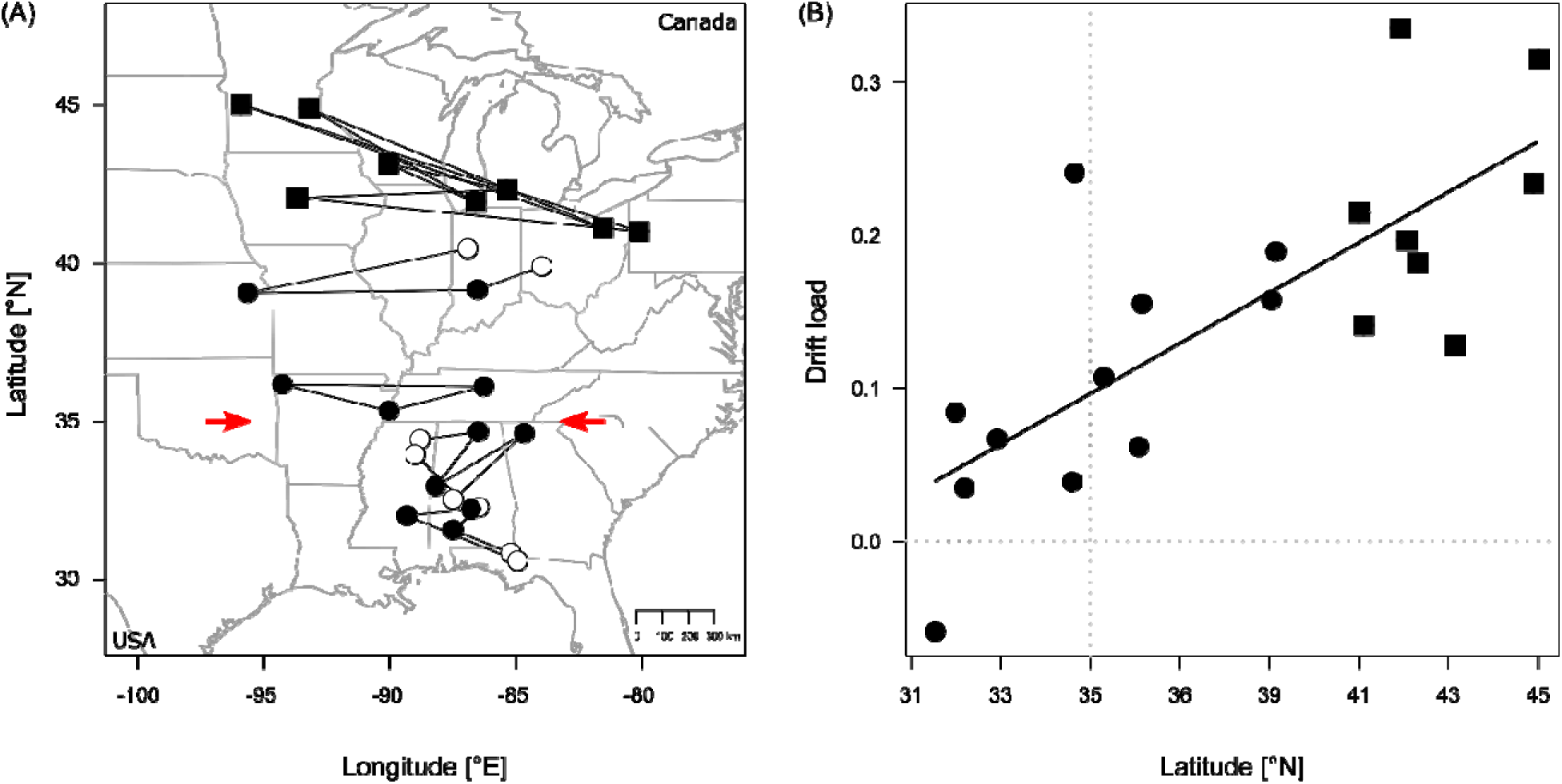
Variation in the expression of drift load across the range (Study 2) **(A)** Location of the populations (dots) crossed (lines) to estimate drift load, with focal populations in black and partner populations in white. Black squares represent additional populations from (Koski et al., 2019). The range of the Western lineage is shaded in light gray, with red arrows representing the latitudinal delimitation of the rear edge. **(B)** Drift load of each focal population (dots) and additional populations (squares) calculated from heterosis estimates of between-population crosses. Values above 0 (horizontal dotted line) indicate drift load expression, values below 0 represent outbreeding depression. The vertical dotted line represents the separation between the rear edge (<35°N) and the expanded range. The solid line represents the significant relationship between drift load and latitude, with the 95% CI indicated in gray shading. Test statistics are reported in Table 1.

For each population, we performed non-reciprocal WPCs among unrelated individuals. For BPCs, 11 populations were chosen as “focal” populations, and were paired with two other populations, either another focal population or one of the 8 remaining populations (Table S2B). Each focal population served as pollen recipient (mother) in one pair, and as pollen donor (father) in the other pair. This design ensured two measures of heterosis per focal population. Pairs were assigned across similar latitudes to minimize environmental differences (Fig. 2A). Hand-pollinations generated 19 WPC lines and 15 BPC lines, with an average 13 seed families per line (Table S2B, Method S2).

We raised on average 21 seedlings for each line (Table S2) between fall 2022 and summer 2023 under similar conditions as described above (Method S2), resulting in a total of 428 WPC and 296 BPC offsprings. Over the course of the experiment, we tracked germination, survival, whether plants flowered, and reproductive output (Method S2, data available in Perrier et al., 2025b).

#### Estimation of drift load

We calculated lifetime fitness at the level of the pot as the product of germination proportion and reproductive output, set to 0 if the plant died or did not flower. Lifetime fitness was averaged across replicates of the seed family within each WPC or BPC line, and then at the level of the line resulting in a single mean for each WPC and BPC line. Heterosis (H) was calculated for each BPC line as the lifetime fitness *(W)* of the BPC line compared to the average lifetime fitness (W) of the WPC of its two parental populations standardized by the average of the parental populations: *H* = (*W*_BPC_ – W _WPC_) / W _WPC_ (Table S2B). Drift load for each focal population was estimated as the average heterosis across its two BPC lines (Table S2A). We supplemented this data with drift load estimates for the additional eight high-latitude populations from Koski et al., (2019; Table S2A).

Based on this combined dataset, we assessed whether rear-edge populations express higher drift load than populations in the expanded range by testing the effect of a populations’ latitude on drift load in a linear model, assuming a normal distribution. Preliminary analysis tested whether latitude was best described by a linear or a second-degree quadratic relationship and results are reported for the best model.

Populations crossed at high latitudes were on average further apart from one another than those at low latitudes, which could potentially affect patterns of drift load due to greater isolation-by-distance and thus differentiation. We therefore explored whether population-average linearized *F*_ST_ (Rousset, 1997) is associated with estimates of drift load in a linear model. For the eight high-latitude populations, we used population-average *F*_ST_ estimates generated from a similar RAD-Seq dataset in Koski et al., (2022).

### Study 3: Do rear-edge populations show strong local adaptation?

We tested the hypothesis that due to a long history of selection under past warming, rear-edge populations exhibit stronger adaptation to local climates than the rest of the range by performing a large common garden experiment. This experiment involved sites representing rear-edge and range-center environments, and tested for fitness advantage of local populations (Kawecki & Ebert, 2004).

#### Experimental design and recording traits

We selected 23 central and rear-edge populations (Fig. 3A, Table S3) to be raised in three common gardens (CG, Fig. 3A, Method S3). Gardens were located along a 850 km latitudinal gradient, representing conditions in the center of the range (CG-Center, Lexington KY, 38.1°N), the rear edge (CG-Rear, Tallahassee FL, 30.5°N), as well as intermediate conditions in between the two (CG-Int, Clemson SC, 34.7°N). This study was also used to assess variation in phenology (Perrier et al., 2025a).

**Figure 3:**
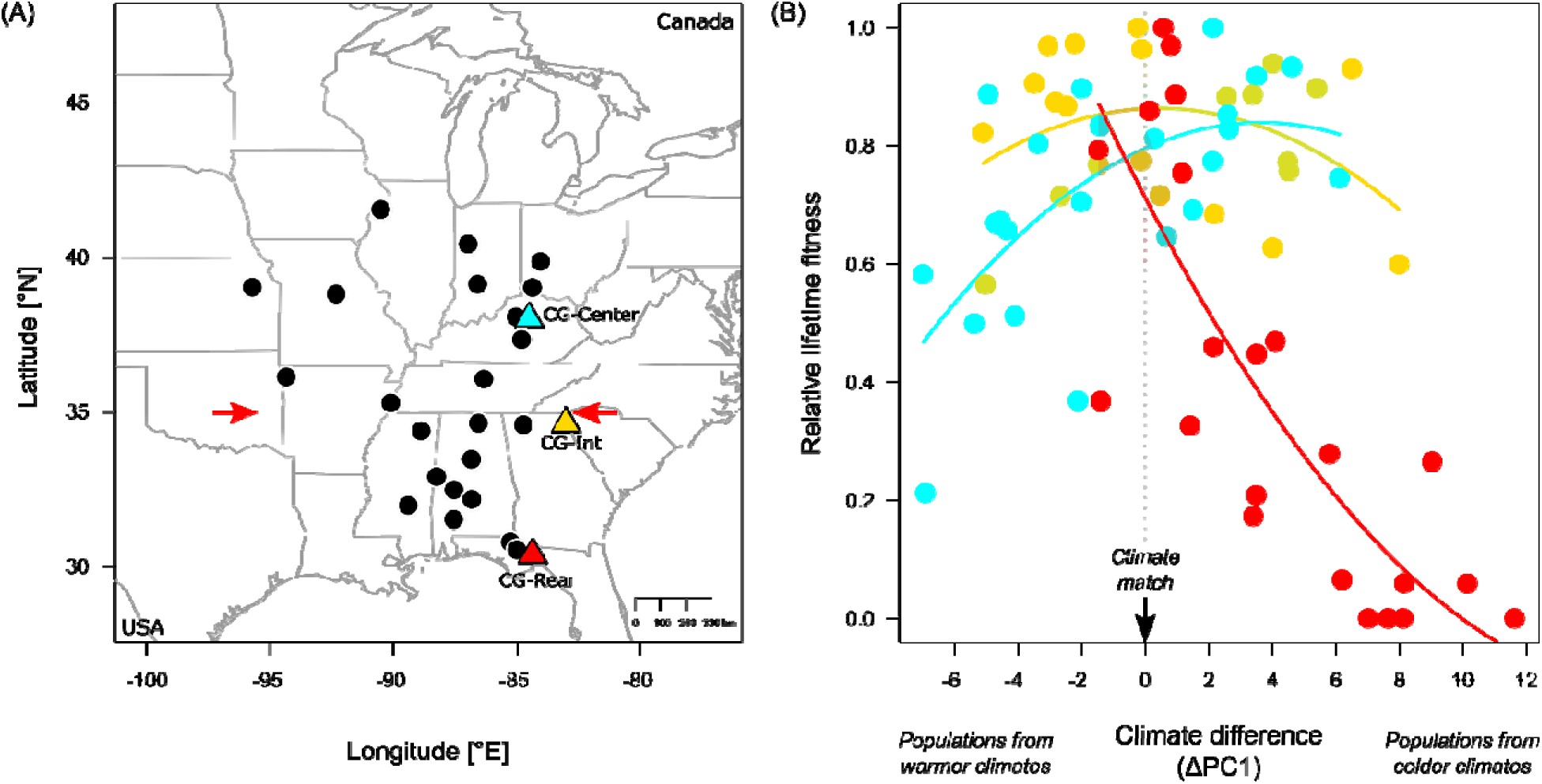
Variation in local adaptation across the range (Study 3) **(A)** Sampling of populations (dots) and common gardens (triangles), with the range of the Western lineage shaded (light gray). Red arrows represent the latitudinal delimitation of the rear edge. **(B)** Relative lifetime fitness of populations (dots) raised in each common garden (distinguished by color as in A). Solid lines represent the relationship between relative lifetim fitness and the climate difference between each garden and population’s location of origin (ΔPC1), estimated for each garden, with the 95% CI indicated in shading (test statistics: Table 2, S7). The dashed line (climate difference = 0) indicates a climate match between the population’s origin and the garden.

**Table 2:** Test of variation in local adaptation (Study 3) across the range of *Campanula americana*.

| Fixed effects | $\chi^2$ |
| --- | --- |
| Common garden (CG) | **6.02** * |
| Climate difference ($\Delta PC1$) | **10.75** ** |
| CG * $\Delta PC1$ | **18.17** ** |

Plants from each of the 23 populations were raised in two cohorts in each common garden (Method S3). The “winter cohort” captured how winter conditions affect survival and the cuing of reproduction, while the “summer cohort” captured the effects of spring and summer conditions on survival and reproduction for seedlings that were vernalized in controlled conditions thereby receiving uniform cuing of reproduction across gardens. For each cohort and garden, we raised 30 seedlings per population (2 individuals/family × 15 families/population) for a total of 4140 individuals (Table S3).

We raised the summer cohort from fall 2022 to late summer 2023. Plants were germinated and vernalized in controlled conditions, transplanted in the field in spring, and harvested in late summer, when most had mature fruits (Method S3). The winter cohort was initiated in fall 2023 and followed through late spring 2024. Seeds were germinated in common conditions and then transplanted in each garden before the onset of winter (Method S3). Data collection finished late spring to allow sufficient time for all plants (that were successfully cued) to bolt after winter. Garden locations were selected to mimic natural habitats of the species (Method S3; Perrier et al., 2025a). For each cohort, one data logger (iButton®; Maxim Integrated Products Inc., San Jose, CA, USA) placed in each garden in the shade, 1.5 m aboveground, monitored air temperature hourly.

We recorded traits across the life cycle (minus germination) for each individual by visiting the gardens multiple times in summer 2023 for the summer cohort, and in spring and early summer 2024 for the winter cohort (Method S3, data available in Perrier et al., 2025b). At each visit, we assessed survival and transition to bolting, and for the summer cohort, transition to flowering (here production of at least one bud). Mature plants of the summer cohort were collected to record reproductive output (total number of fruits, flowers and buds; Method S3).

#### Climate difference between common gardens and populations

To quantify climate differences between the conditions experienced in the common gardens and each population’s location of origin, we generated 19 temperature and precipitation related variables that captured seasonal and life-stage-specific variation (Table S4, Method S3). For gardens, climate data came from dataloggers and nearby weather stations during the experiment; for populations, from the PRISM database (2018–2022, https://prism.oregonstate.edu, accessed 13/05/2024). To reduce dimensionality and autocorrelation, we performed a PCA on the population climate variables. Climate variation was mostly explained by an axis summarizing variation in winter and spring temperature (PC1, 76.65% of variance explained, Table S5), and by an axis summarizing variation in precipitation and summer temperatures (PC2, 8.81% of variance explained, Table S5). The gardens were projected into this PC space and captured the range of variation along PC1 but less variation along PC2 (Figure S1). We thus focused on PC1 to assess local adaptation. For each population in each garden, we calculated the climate difference (ΔPC1) between the garden’s PC1 and the population’s home PC1 (ΔPC1 = PC1_garden_ – PC1_population_), where a value of 0 indicates climate matching, i.e. the population’s climate is “local” to the garden, while positive or negative values indicate that populations experience warmer or colder conditions in the garden than at their location of origin.

#### Assessment of local adaptation

We generated a measure of lifetime fitness across the two cohorts by combining five traits summarizing fitness at key life stages: survival over winter, survival until reproduction, bolting, flowering and reproductive output (Method S3). Traits were averaged at the seed family level in each garden. For a few family–common garden combinations, data on traits in either cohort was missing (summer cohort: 20 of 992, winter cohort 43 of 1015). For these cases, missing trait data was imputed as the average across populations in the garden. Family averages were then multiplied across life stages to estimate lifetime fitness. Fitness was then calculated for each population within each garden after log10 transformation (+1 so that all values are > 0) to achieve normality. Population-level lifetime fitness was standardized to account for variation in garden quality by dividing each value by the maximum value within that garden.

To evaluate whether local adaptation is stronger at the rear edge, we tested how variance in fitness depended on the climate difference between the populations and gardens. If populations are locally adapted, we expect fitness in a given garden to be highest for populations where the climate at their location of origin matches the garden’s climate (ΔPC1 = 0), and to decline with increasing climate difference (*i.e.* following a local vs foreign framework, Kawecki & Ebert 2004). Stronger declines in fitness indicate stronger local adaptation. Variation in relative lifetime fitness was tested in a hierarchical mixed effect model (Kuznetsova et al., 2017), assuming a normal distribution. Fixed effects included common garden (three levels), climate difference (ΔPC1), and their interaction. The effect of climate difference was modeled using a quadratic relationship to allow fitness to decline towards positive and negative values of ΔPC1. Population was included as a random effect. Post hoc tests (Lenth, 2019) evaluated differences in fitness among gardens, and in the climate-fitness relationship between gardens. Finally, we tested the extent to which variation in population-level lifetime fitness was explained by a population’s latitude as fixed effect.

## Results

### Study 1: Are rear edges reservoirs of diversity?

Rear-edge populations are less genetically diverse but more genetically differentiated than central populations. Patterns of genetic diversity within populations and differentiation across populations were best explained by a quadratic relationship with latitude (Table S6). Diversity based on θ_W_ was highest for mid-latitude populations and significantly declined at high and low latitudes Table 1, Fig. 1B). While not significant, θ_π_ followed a similar pattern (Table 1, Fig. S2). In contrast, differentiation between populations, accounting for geographic distance, was lowest at midlatitudes and significantly increased towards low latitudes, and to a lesser extent towards high latitudes (Table 1, Fig. 1C).

### Study 2: Do rear-edge populations suffer from strong genetic drift?

Rear-edge populations expressed less drift load than the expanded range. Most populations across the range displayed some load (Fig. 4B), indicated by positive values. Variation in drift load was best explained using a linear model (Table S6), and increased significantly with latitude (Table 1, Fig. 2B). There was no relationship between drift load and linearized F*_ST_* (*F* = 0.01, *P*=0.93, *R*^2^ = 0.00, Fig. S3), indicating that isolation-by-distance (Fig. 2A) did not drive variation in drift load.

### Study 3: Do rear-edge populations show strong local adaptation?

Local adaptation, while common across the range, is highest at the rear edge. In each garden, climate-matching populations (i.e. local) generally had higher fitness than populations from warmer or colder climates than experienced at the garden (Table 2, Table S7B). The strength of local adaptation (slope of climate difference on fitness) was similar in the central and intermediate gardens, but stronger in the rear edge garden (significant interaction, Table 2, Table S7C), where populations from increasingly colder climates had increasingly lower fitness, with the most northern populations having a fitness of zero (Fig. 3B).

The analysis of lifetime fitness across latitudes yielded similar results (Table S8). Fitness in each garden was highest for populations originating from similar latitudes as the garden (Fig. S4). The highest fitness overall was achieved by low-latitude rear-edge populations in the rear-edge garden.

## Discussion

Rear-edge populations, relicts that persist in glacial refugia, offer unique insights into how populations evolve under climate change. We investigated evolutionary responses to past warming in the North American plant *Campanula americana* by comparing genetic patterns between populations at the rear edge and in the expanded range. Rear-edge populations exhibited reduced genetic diversity and higher differentiation among populations, typically interpreted as evidence of genetic drift, yet show minimal drift load. Instead, rear-edge populations expressed strong local adaptation, suggesting they have persisted in place by gradually adapting to changing climates. Our results highlight the importance of considering and testing multiple evolutionary responses to past climate change when inferring historical processes from contemporary genetic patterns. Our findings also identify local adaptation as a key response to past (and potentially future) warming, and reveal rear edges as powerful models of successful adaptation under long-term climate change.

### Estimates of genetic variation and drift load support contrasting evolutionary histories

Past climate warming left mixed signatures of evolutionary processes at the rear edge of *C. americana.* Estimates of genetic variation do not support the hypothesis that these populations act as reservoirs of genetic diversity, as expected for stable rear edges (Perrier et al., *in review*). Instead, they point toward strong genetic drift with lower within-population diversity at the rear edge than in central populations, particularly for segregating alleles (θ_W_), indicating a depletion of rare alleles. However, rear-edge populations also showed markedly higher differentiation among populations compared to the rest of the range, including the leading edge. Diversity between populations rather than within them is often attributed to strong genetic drift, especially at the rear edge (e.g. Scalfi et al., 2009; Diekmann & Serrão 2012; Assis et al., 2014; Carbognani et al., 2019; Kebaïli et al., 2022; but see Wood et al., 2021).

Previous work in the species provides ecological context for these patterns. Habitats at *C. americana’s* rear edge served as refugia, and were highly suitable for the species during the LGM. However, they are now of low suitability (Barnard-Kubow et al., 2015), likely reflecting a long history of habitat degradation due to warming since the LGM. Habitat degradation is expected to lead to demographic decline, progressive isolation, and a reduction in diversity in populations. We found within-population diversity also declined toward high latitudes, perhaps reflecting cumulative genetic erosion of populations established through repeated bottlenecks during postglacial range expansion (Excoffier et al., 2009). Together, these patterns suggest that past climate warming led to reduced diversity at both range edges, but likely through different mechanisms, with expansion-driven erosion at the leading edge and long-term habitat decline at the rear edge.

Contrary to expectations, rear-edge populations expressed the lowest drift load across *C. americana*’s range. Therefore, results do not support the hypothesis that these populations have experienced strong genetic drift. Further, the results contradict expectations of drift inferred from within- and between-population estimates of genetic variation (Study 1) and habitat decline at the rear edge (Barnard-Kubow et al., 2015). Instead, drift load increased toward high latitudes, consistent with the decline in genetic diversity observed at the leading edge and the expectation that drift increases with range expansion (Peischl et al., 2013; 2015). Our findings contrast with the only previous evaluation of drift load at the rear edge that found it elevated and of similar magnitude to the leading edge in *Arabidopsis lyrata* (Willi et al., 2018; Perrier et al., 2020). Overall, while patterns of genetic diversity and drift load suggest that past warming led to an expansion-driven history of drift at the leading edge, other evolutionary processes – beyond genetic drift – appear to have shaped genetic diversity and differentiation at the rear edge.

### Strong local adaptation allowed rear-edge populations to persist in declining habitats

In the common garden experiment, populations generally performed better in climates similar to conditions at their location of origin, consistent with local adaptation (Kawecki & Ebert, 2004; Leimu & Fischer, 2008). This pattern was strongest in the rear-edge garden, where the fitness of populations declined sharply as climates became increasingly different from a population’s home. Strong local adaptation at the rear edge likely reflects a history of persistent directional selection under ∼20kya of postglacial warming, confirming the third hypothesis. This long-term environmental change likely provided the selective pressure and time required for gradual adaptation to increasingly warm conditions, enabling rear-edge populations to persist in place despite experiencing drastic changes in habitats. Local adaptation was weaker further north where populations tracked shifts in suitable habitats and therefore experienced less environmental change, providing less opportunity for local adaptation than in the rear edge. Elevated local adaptation towards the equatorial range limit is common across temperate species (Bontrager et al., 2021). Our findings experimentally corroborate this pattern and demonstrate strong local adaptation as a potential hallmark of rear edges.

Rear-edge populations in *C. americana* thrived in their local garden despite environmental conditions that appear marginal for the species. In the rear-edge garden, populations from the expanded range had very low fitness, sometimes as low as zero, primarily due to a failure to initiate reproduction (Perrier et al., unpublished data). Lack of reproductive cuing was likely due to insufficient winter cold. Most populations of the species require much longer periods of vernalization for successful cuing (Perrier et al., 2025a) than the brief two days experienced in this garden (Table S4), though environmental conditions in this garden are typical of the subtropical climates found at the rear edge (Perrier et al., 2025a). Contemporary rear-edge habitats may thus fall outside of the climate niche occupied by most of the species as winters are too warm (Perrier et al., 2025a). This finding is consistent with the low suitability at the rear edge of the species found by niche modeling (Barnard-Kubow et al., 2015), and reflects general expectations of the marginality of rear edge habitats (Bontrager et al., 2021; Vilà-Cabrera et al., 2019). Yet, plants from rear-edge populations did not suffer from the low quality habitats in the rear-edge garden, and in fact achieved some of the highest performance across populations and gardens. Our findings contribute to growing evidence that rear edges may harbor unique adaptations to warming (Ghouil et al., 2020; Pelletier et al., 2023; Perrier et al., 2025a), allowing them to thrive in distinct warmer habitats that are unsuitable relative to the niche of the species as a whole. These findings also echo recent critiques of predictions of marginality at the rear edge (Vilà-Cabrera et al., 2019), which often do not take local adaptation into account. As we demonstrate here, overlooking local adaptation may skew expectations of performance at range limits.

### Revisiting expectations of evolution under warming climates

Our study in *C. americana* offers new insight into how past climate change may have shaped rear-edge populations. Most studies on the rear edge evaluate one of two possible evolutionary trajectories – high levels of ancestral diversity or drift-driven decline (Perrier et al., *in review*). Our findings demonstrate a third, often overlooked, possibility that rear-edge populations can persist and even thrive under long-term climate change, not because they maintain high genetic diversity, but because they have gradually adapted to changing conditions. This third evolutionary trajectory offers a more nuanced framework for understanding dynamics under past climate change, and spotlights rear edges as models for studies of how populations may successfully adapt to continuously warming climates.

Our study also challenges the assumption that reduced diversity and heightened differentiation are reliable evidence for strong genetic drift. This pattern has been found in many studies, contributing to the prevalent notion that rear edges experience strong drift (Perrier et al., *in review*). Yet, our results indicate that this assumption does not always hold, and suggest other evolutionary forces may create this pattern. For example, historic selection linked to strong local adaptation could have driven selective sweeps reducing diversity (De La Torre et al., 2021; Wei et al., 2023). Likewise, high differentiation of populations at the rear edge might simply be the result of the long history of populations in this area. Only a few studies have gone beyond estimates of genetic variation to test for drift, i.e. genetic signatures of population decline (Rodríguez-Muñoz et al., 2007; Cho et al., 2020; Dupoué et al., 2021; Kvist et al., 2015; Lepais et al., 2022), or drift load (Perrier et al., 2020, 2022). One of those suggested that low diversity and high differentiation in seaweed may result from past selection, not drift (Wood et al., 2021). The lack of explicit testing, and the fact that local adaptation can hide in plain sight, masquerading as drift, raises the question of how often rear-edge populations thought to have been molded by drift are actually shaped by historical selection and adaptation? Resolving this question will require broader consideration of evolutionary histories at the rear edge and targeted tests to distinguish them.

### Evolutionary legacies and the fate of rear-edge populations under future climates

Past evolutionary processes may influence how rear-edge populations respond to future climate change. For example, rear-edge populations are often expected to experience high drift (Hampe & Petit 2005; Perrier et al., *in review*), raising widespread concerns about their long-term persistence (Nadeau & Urban, 2019; Vilà-Cabrera et al., 2019). In small range-edge populations, drift can indeed reduce adaptive potential, reduce fitness, and increase vulnerability to warming stress (Koski et al., 2019; Perrier et al., 2020, 2022; Sánchez-Castro et al., 2022). As we show here, this vulnerability at the rear edge may be overestimated, because drift is often inferred from estimates of genetic variation alone. Our findings of strong local adaptation at the rear edge also raise questions about the long-term impacts of this evolutionary legacy, as local adaptation can have contrasting consequences for populations under changing environments (Chai et al., 2025). On one hand, adaptation to past warming could improve populations’ responses to expected future warming by granting higher resilience and facilitating future adaptation. For example, in *C. americana*, rear-edge populations are expected to be more resilient to future warmer winters than more northern populations due to their reduced vernalization requirements (Perrier et al., 2025a). On the other hand, the same evolutionary trajectory may come at a cost if strong historic selection leads to reduced genetic diversity which limits standing genetic variation necessary for future adaptation (Barrett & Schluter, 2008). Moreover, specialization to current local climates may increase sensitivity to environmental shifts, leading to increasing risk of maladaptation under future climates (Frank et al., 2017; Kooyers et al., 2019; Anderson & Wadgymar, 2020). In sum, the evolutionary legacies in rear-edge populations need to be further characterized to inform robust evaluations of resilience and vulnerability at warmer range limits.

## Conclusion

Our study in *C. americana* highlights rear-edge populations as powerful models to understand how populations evolve under long-term climate change. Realizing this potential requires the integration of genomic and experimental approaches as well as the consideration of multiple possible evolutionary trajectories. More broadly, our study suggests that rear-edge populations are not necessarily relics in decline but products of successful, gradual adaptation to warming climates. As such, they represent valuable yet underappreciated natural laboratories to study the mechanisms enabling persistence under changing climates. Recognizing them as models of successful adaptation – not just vulnerability – could provide valuable insight for forecasting and managing species responses under future warming.

## Acknowledgements

This work was supported by the Swiss National Foundation (P2BSP3_195363) and the University of Virginia College of Arts and Sciences. We are grateful to David Westneat (University of Kentucky Ecological Research and Education Center Field Station, Lexington, KY), Trevor Stamey (Clemson University Experimental Forest, Clemson, SC), Theresa Jepsen (Florida State University Mission Road Research Facility, Tallahassee, FL) for logistical support in establishing common gardens. For their help in raising of plants, performing crosses, setting up the common garden experiments, and collecting data, we thank A. Burricks, C. Claussen, S. Cox, L. Elhady, M. Gower-Fici, K. Haines, S. Kelly, K. Lamb, A. López, H. Makowski, M. Marcich, L. Pizarro, E. Scott, E. Savier, & M. Turner. We are also grateful to D. Brown, J. Collins, A. Diamond, L. Elliott, E. Galloway, F. Griffith, I. Guenther, J. Hansen, H. Horne, J. Kees, M. Kohout, R. Laporte, D. Reed and B. Sutherland for seed and leaf material collection in natural populations. Collection permits were provided by the Florida Department of Environmental Protection, the Tennessee Department of Environment and Conservation, and Missouri Department of Conservation.

## Conflict of Interest Statement

The authors declare that there is no conflict of interest.

## Author Contributions

All authors contributed to the study design. AP collected seeds in the field, prepared samples for sequencing, and performed crosses. AP and OK raised plants and collected data in the greenhouse experiment. AP raised plants and collected data in the field experiment. JB helped in designing the sequencing experiment and with the analysis of genetic data. AP analyzed the genetic and phenotypic data. AP wrote the manuscript with input from LG and JB.

## Data availability statement

Raw read data are publicly available in the Sequence Read Archive (BioProject PRJNA1306192). Custom scripts necessary to recapitulate our genetic analyses, as well as the diploid VCF, loci-level summary statistics, pairwise *F*_ST_ and geographic distance, fitness data of the drift load and local adaptation studies, are stored on Zenodo (https://doi.org/10.5281/zenodo.16883804).

## Notes

### Competing Interest Statement

The authors have declared no competing interest.

