## Supplementary Methods for "The legacy of past climate warming: strong local adaptation in rear-edge populations"

### Method S1: Population genetic study (Study 1)

*Plant material*

We collected leaf material from seven to ten individuals of 31 populations, for a total of 292 individuals (Fig. 1, Table S1.1). Leaves were collected on individuals raised in the greenhouse from seeds harvested from presumably unrelated maternal plants in natural populations (107 samples) or from greenhouse produced seeds (independent within-population crosses, 185 samples, see Method S2). Samples were gently dried in an oven at low heat (~35 to 40 °C) for two to three days, then preserved at -20 °C until sequencing.

*DNA extraction, library preparation & sequencing*

Samples were sequenced using a Restriction-site Associated DNA sequencing approach (RAD-Seq; Andrews et al. 2016; Baird et al. 2008). These samples were part of a larger sequencing project of 475 individuals (Bioproject PRJNA1306192). DNA extraction was done by the Genomics & Cell Characterization Core Facility (GC3F) of the University of Oregon (Eugene, OR, USA). For each sample, DNA was extracted from at least 20mg of dry leaf tissue (MagMAX™ Plant DNA Isolation Kit, Applied Biosystems, Foster City, CA, USA) then purified (Mag-Bind® TotalPure NGS, Omega Bio-tek, Norcross, GA, USA). Samples were then concentrated, tested for purity and fragmentation, and equilibrated to 20 ng/µL.

Library preparation and sequencing was performed by Floragenex, INC (Beaverton, OR). Samples were processed in five batches of 95 individuals from extraction to sequencing. Individual samples were digested with the *PstI* restriction endonuclease, and resulting fragments were ligated with adaptors containing unique 10 bp barcodes to identify each sample. Fragments were pooled by groups of 95 (multiplexed), sheared by sonication, size selected (300–500 bp range) and amplified by PCR. Libraries were paired-end sequenced (150bp) on two S4 flow cell lanes per library using a NovaSeq6000 (Illumina, Inc. San Diego, CA, USA). For two of the five libraries, initial sequencing had low yields. These libraries were sequenced two additional times.

*Processing of raw reads*

Raw reads were processed using a modified STACKS v2 pipeline for RAD-Seq data using a reference genome (Rochette et al., 2019). Multiplexed reads were first quality-checked using *fastqc* (Andrews, 2010). For each sequencing lane and library, paired reads were demultiplexed (assembled by individual), filtered for low quality and missing bases (-c -q) and trimmed to remove barcodes using *process-radtags* in STACKS. For each individual within each lane, PCR duplicates were removed using *clone_filter* in STACKS. The remaining sequence data was then merged within individuals across sequencing lanes. For each individual, forward and matching paired-end reads were aligned to a *C. americana* reference genome (haploid chromosome-level assembly based on the Appalachian clade population VA73 using PacBio HiFi reads; A. Lopez-Caamal, *unpublished data*) using BWA-MEM (Li, 2013; Li & Durbin, 2009). Based on these alignments, we generated a catalog of RAD loci (groups of homologous reads derived from the same genomic region and assembled across samples) including all individuals, using *gstacks* in STACKS. In total, the catalog comprised 1,566,971 RAD loci, assembled from 924,427,374 forward reads and 748,416,814 matched paired-end reads, for an average coverage of 31.2x per individual (SD = 9.7x; range: 8.8x – 80.6x). RAD loci averaged 465bp in length with a mean insert length of 72bp, for a total length of 615 Mbp, covering about ~36% of the 1.7 Gbp haploid reference genome (A. Lopez-Caamal, *unpublished data*).

*Genotyping*

Genotypes were called from the catalog generated above as variant call format (VCF) using *populations* in STACKS. We retained SNPs where sequence information was present for at least half of the individuals within each population (-r 0.5, not taking account populations where the site was entirely missing) and for 85% of individuals overall (-R 0.85), and with a minor allele count of at least 2 (--min-mac 2) to include rare variants for accurately calculating diversity estimates while removing singletons that could result from errors during sequencing. The resulting 158,356 SNPs were filtered to exclude 16,778 SNPs with a mean read depths of less than 20 (following Koski et al. 2019) or greater than 67 (average coverage + 4*√average coverage, following (Li, 2014); average before filtering = 41) using VCFTools. In addition, 4,266 variants resulting from indels were excluded using BCFTools (Danecek et al., 2021). This resulted in a final set of 137,312 high quality SNPs (average coverage of 38.9x reads per individual) distributed across 7,602 RAD loci, which together span ~7.3 Mbp, corresponding to ~0.4% of the genome sequenced in each individual. For each sample and at each SNP, diploid genotypes were converted into tetraploid genotypes to account for partial heterozygotes. Tetraploid genotypes were estimated using a custom R script by first calculating the ratio of reads supporting the derived allele (B) over the reference allele (A) from the number of reads provided in the VCF file, then assigning genotypes using following bins: ≤ 0.05, AAAA; <0.375, AAAB; <0.625, AABB; <0.95, ABBB; ≥ 0.95, BBBB. This method was used in a previous study in the species to estimate tetraploid genotypes from RAD-Seq data (Prior et al., 2020), with the difference that we added the thresholds of 0.05 and 0.95 to bin homozygous genotypes (instead of 0 = AAAA and 1 = BBBB), to account for potential sequencing errors. These thresholds represent cases where out of the minimum of twenty reads necessary to retain a SNP, only one read supports the minor allele.

*Accounting for isolation by distance in measures of genetic differentiation among populations*

Because sampling across the range was not homogeneous and genetic differentiation among populations is often expected to increase with geographic distance due to isolation-by-distance (IBD; Eckert et al. 2008; Slatkin 1993), we accounted for IBD before assessing variation in differentiation across the range. We estimated differentiation by calculating pairwise $\mathrm{F}_{\mathrm{ST}}$ (Weir & Cockerham, 1984) among populations using the *StAMPP* package in R (Pembleton et al., 2013). We also estimated geodesic geographic distance (km) for each pair of populations using the harvestine method implemented in the *geodist* package in R (Karney, 2013). Pairwise $F_{\mathrm{ST}}$ estimates were linearized ($F_{\mathrm{ST}}$ / (1- $F_{\mathrm{ST}}$ ) to account for the non-linear relationship between $F_{\mathrm{ST}}$ and geographic distance under isolation-by-distance models (Rousset, 1997), and included in a linear model testing for the effect of pairwise geographic distances ($R^2$= 0.08, P < 2e-16). Residuals of pairwise $\mathrm{F}_{\mathrm{ST}}$ were then averaged at the level of the population as a measure of how differentiated a population was from others relative to the expected differentiation based on IBD (Table S1).

**Method S2: Drift load study (Study 2)**

*Plant growth in greenhouse conditions*

Seeds of different maternal plants (hereafter seed families) were collected in late summer 2020 and 2021 in natural populations and stored in separate bags at 4°C under dark and dry conditions. We sowed up to five seeds for each seed family in each population in each of two pots filled with a 3:1 mixture of potting mix (LM-111 All Purpose Mix, Lambert, Rivière-Ouelle QC, Canada) and turface. Pots were randomly distributed across 128-cell propagation trays. Germination occurred in growth chambers under 12h light, 21°C day, 14°C night, 60% RH for four weeks. Pots were watered daily for a week, then every other day. Trays were randomized within growth chambers twice a week. At the end of the germination, seedlings were randomly thinned to one per pot and plants were fertilized once (13 mg/L, Jack's Professional 15-5-15 + CaMg LX Fertilizer, JR Peters Inc., Allentown PA). Seedlings were then vernalized to simulate winter (4°C, low light for 12h per day) for six weeks, which is required for most populations to initiate reproduction (Perrier et al., 2025).

After vernalization, one replicate per seed family was retained and moved to a greenhouse with daylength extended to 16h with artificial light (600 µmol/m2/s) to induce flowering. Initial temperature of 23°C day / 19°C night (+/- 5°C) was increased after two months to daytime temperature of 25°C and plants were shaded to simulate a light environment of an open forest. These temperatures reflected average summer temperatures across the range (Table S4). Plants were watered as needed. Fertilizer was applied at 1000ppm (N) every two days for two months, then one to two times a week. Plants were treated with fungicide (two times before and two times after vernalization, Banrot 40 WP, Everris NA Inc., Dublin, OH; 0.9 g/L) to avoid root rot. During flowering, plants were sprayed with pesticide once a week to reduce damage by thrips (neem oil and liquid soap 1:1, diluted in water at 13ml/L, and spinosad diluted in water at 16.5ml/L).

*Generation of within- and between- population crosses*

Each plant within a population served as pollen recipient (mother) and was paired with two randomly selected other individuals of the same population to serve as pollen donors (fathers). This resulted in two within population cross-combinations (WPC) per plant (i.e. seed family), with each plant serving as mother and as father twice at most. Between-population crosses (BPC) were performed by selecting 11 "focal" populations, and pairing each with two other populations out of the 19 available (Fig. 2), to ensure two independent measures of heterosis per focal population. Pairs were assembled at roughly similar latitudes to minimize the effects of divergent local adaptation. In this design, each focal population served as the mother population in one pair, and the father population in the other pair. For 7 focal populations, both pairs involved other focal populations. In total, we generated 15 BPC lines (Table S2B). For each BPC line, 15 plants of the mother population (that were also mothers in WPC crosses) received pollen from one randomly chosen plant from the father population.

All crosses were performed by hand on emasculated flowers on the mother plants with pollen from a flower collected on the selected father plant. Each cross-combination was repeated on three female flowers for WPC, two for BPC. If all repetitions of a cross combination failed, the father plant was changed. Mature fruits were collected and stored in individual envelopes at 4°C under dry and dark conditions.

*Raising of WPC and BPC offspring*

We grew 18 to 25 individuals for each WPC and 18 to 20 for BPC, with each seed family represented twice but occasionally once or three times, depending on how many seeds were available per family (Table S2). For each pot, we sowed two seeds and thinned to one individual before vernalization. Plants were raised as described above, except that germination lasted five weeks and vernalization lasted seven weeks. This experiment was stopped once all plants that successfully bolted had flowered for five weeks

*Recording of fitness traits*

We tracked fitness across the life cycle. We scored germination and survival of each seedling once a week before simulated winter. After plants were moved to the greenhouse, survival and flowering (opening of at least one flower) were scored three times a week for each plant. Once a plant started flowering, we recorded its reproductive output as the sum of weekly open flower counts over five weeks.

**Method S3: Local adaptation study (Study 3)**

*Raising of plants in the common garden*

Plants of the 23 populations selected for the local adaptation study were raised in three common gardens in two cohorts (Method S3 Figure A, B). We used greenhouse-reared seeds (WPC, Method S2) for 20 populations, and field collected seeds for the remaining three (Table S3). For each population, we selected 15 seed families (randomly duplicated if fewer available) to be used in the experiment. With few exceptions (1 families out of 345 in the first cohort, 3 out of 347 in the second), we used the same seed families in both cohorts.

We raised the summer cohort from fall 2022 through late summer 2023 (Perrier et al., 2025). For each garden, two seeds per family (three for field seeds, one if few seeds) were sowed in each of two pots, thinned to one per pot after germination. Seeds were germinated for 6 weeks and vernalized for 7 weeks as described in Method S2. In spring 2023, seedlings were acclimated outdoors for 1 week (Charlottesville, VA, USA), then transplanted into each CG, starting with the southernmost garden to reflect the earlier start of the growing season at lower latitudes. The transplant date (Method S3 Table A) was chosen as the date when mean daily temperature was consistently above 10°C in spring (estimated using data for 2018–2022 from the PRISM database, https://prism.oregonstate.edu, accessed 13/05/2024, more details in Perrier et al. 2025). Sowing dates were planned to ensure seedlings were transplanted at the same life stage across CGs.

The winter cohort was initiated in fall 2023 and followed through late spring 2024. The seed sowing design was similar to the summer cohort. Seeds were germinated for 5 weeks as in Method S2, then acclimated for four days to the colder outdoor conditions of late fall (Charlottesville, VA), except for the southernmost garden where seedlings were kept in indoors growing conditions that better reflected the temperatures at the garden's location. Seedlings were then transplanted in each garden starting with the northernmost garden, reflecting the earlier start of winter. Transplant date (Method S3 Table B) was chosen as the date two weeks before the first day where mean daily temperature was below 4°C in fall, or two weeks before the first day of frost, depending on which occurred first (dates estimated using PRISM data for 2018–2022). Sowing dates were planned accordingly.

In both cohorts, plants were transplanted into 30 blocks of 23 individuals (Method S3 Figure B), with one individual per population per block resulting in 30 replicates of a population in each garden (final replication in Table S3, more details in Perrier et al., 2025). The distribution of populations within blocks, and of seed families for each population between blocks, was determined though randomization, and was changed between both cohorts. The same garden locations were used for both cohorts.

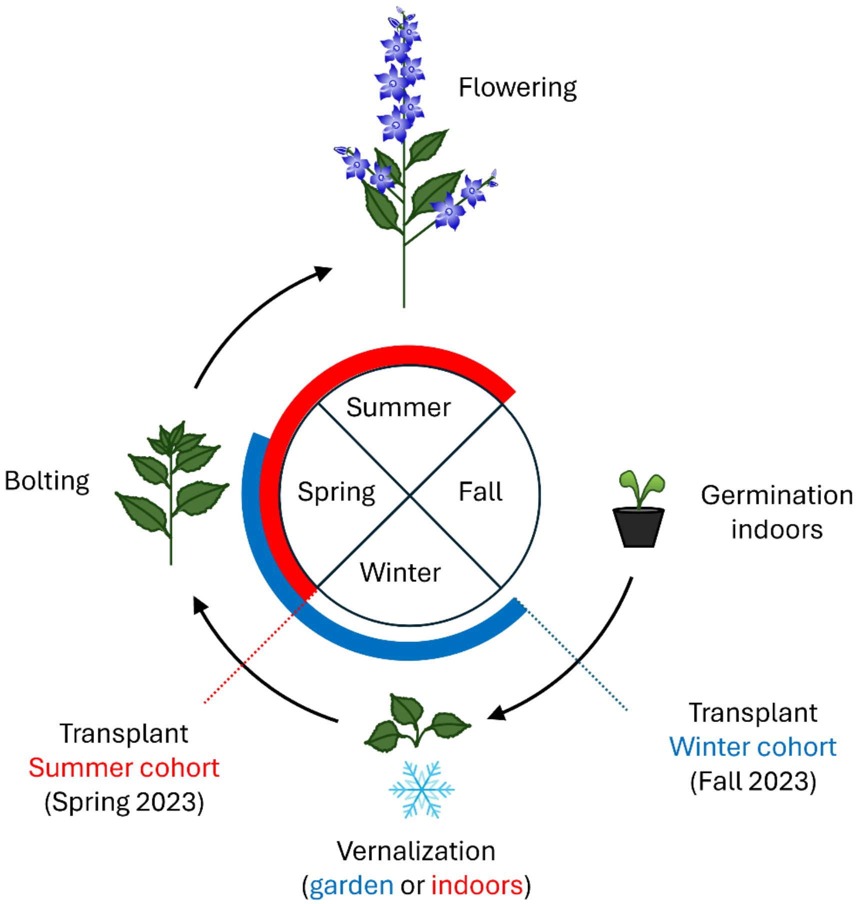

**Method S3 Figure A: Life cycle of *C. americana* split between winter and summer cohorts.** The common garden experiment consisted of a summer cohort (red, spring and summer 2023) and a winter cohort (blue, fall 2023 – spring 2024). Plants were germinated indoors for both cohorts, then transplanted into each garden before winter for the winter cohort, and after a simulated winter indoors for the summer cohort. Together, the two cohorts allowed determining the effects of climate across most of the species' life cycle.

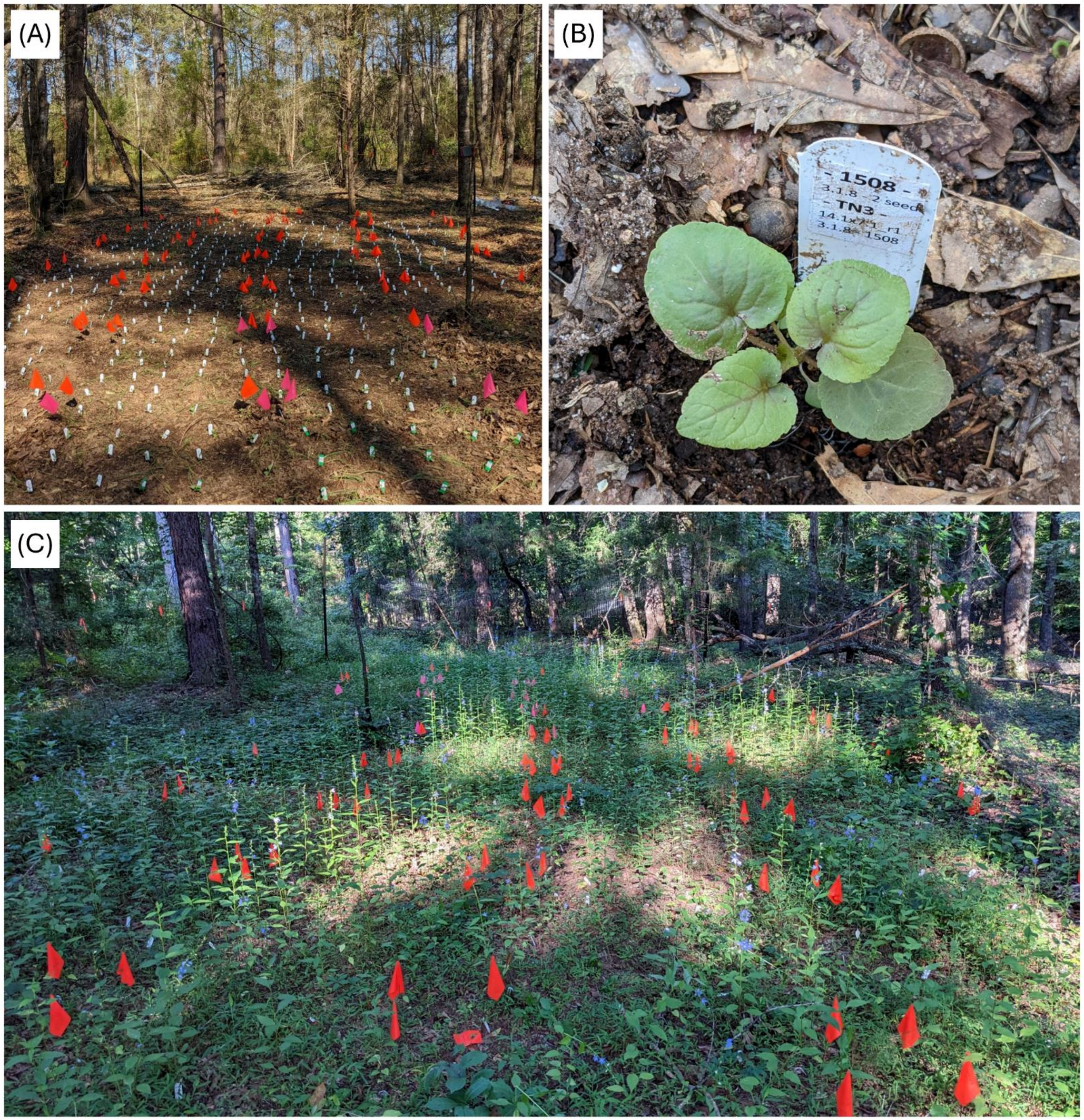

**Method S3 Figure B: Pictures of the common garden CG-Int (Clemson, SC), summer cohort**
**(A,B)** Garden right after transplant. (C) Garden on the first visit in summer.

**Method S3 Table A: Location of common gardens (CG) and schedule of the summer cohort**

| | Latitude | Longitude | Transplant | Visit 1 | Visit 2 | Visit 3 |
|---|---|---|---|---|---|---|
| CG-Core | 38.08079 | -84.47134 | 08/04/2023 | 16/07/2023 | 12/08/2023 | 09/09/2023 |
| CG-Int | 34.68746 | -82.87249 | 22/03/2023 | 14/07/2023 | 13/08/2023 | 06/09/2023 |
| CG-Rear | 30.45723 | -84.33319 | 02/03/2023 | 12/07/2023 | 17/08/2023 | - |

For CG-Rear, 98% of plants were considered mature at the second visit, thus all plants were collected, and the garden was not visited a third time.

**Method S3 Table B: Schedule of the winter cohort**

| | Transplant | Visit 1 | Visit 2 |
|---|---|---|---|
| CG-Core | 29/10/2023 | 28/03/2024 | 05/06/2024 |
| CG-Int | 12/11/2023 | 25/03/2024 | 31/05/2024 |
| CG-Rear | 14/11/2023 | 26/03/2024 | 02/06/2024 |

*Recording of fitness traits*

We defined five traits representing fitness components at different life stages. *Survival over winter* (binary) was the survival of plants in the winter cohort from transplant in fall (2023) until the first recording date in early spring (2024). *Survival until reproduction* (binary) was the survival of plants in the summer cohort from transplant in early spring (2023) until the end of the cohort in summer (2023). *Bolting* (binary) was the successful transition to bolting of surviving plants in the winter cohort and was assessed over two visits (2024). *Flowering* (binary) was the successful transition to flowering of plants in the summer cohort (2023) that had bolted. *Reproductive output* was the sum of fruits, flowers and buds (see below) of plants that successfully flowered in the summer cohort (2023).

*Estimation of reproductive output*

In the summer cohort, mature plants were collected at each visit to determine reproductive output. Plants were considered mature if they had been flowering for at least 5 weeks (first fruits starting to turn brown, see Perrier et al. 2025). This timing was chosen to prevent mature fruits from opening and dispersing seeds. At the southern garden (CG-Rear), all plants were collected at the second visit as 95% of plants had flowered (Perrier et al., 2025), and of these 98% were considered mature. In the other two gardens, all reproductive plants were collected on the third visit, in September, regardless of maturation. In most natural populations, flowering is finished by September (personal obs.), and open flowers at this time are unlikely to produce mature fruits before the onset of winter. Stopping the experiment then thus allows for an accurate representation of the contribution of phenology to reproductive success, an important component of adaptive divergence between populations across the range (Perrier et al., 2025). Upon collection, plants were cut at the base of the flowering stalk, stored in individual bags and dried. We scored reproductive output as the sum of the number of fruits, flowers and buds on each plant. We also measured the stem diameter at the base of the flowering stalk.

For 325 plants (~20% of all plants that reproduced, Method S3 Table C), reproductive output could not be scored accurately. These included plants that matured before collection started and lost fruits, plants where the main stem was broken, and 15 plants that did not reach fruit maturation in the southern garden. For these, reproductive output was predicted based on the relationship between reproductive output and stem diameter. We estimated this relationship across gardens for each population using linear models with reproductive output as the dependent variable (assuming normal distribution), and the squared stem diameter (in mm) as fixed effect (average $R^2 = 0.60$, Method S3 Table C). In cases where the predicted reproductive output was lower than the minimum observed reproductive output for that population, the predicted value was assigned to be the minimum observed value.

**Method S3 Table C: Summary statistics for reproductive output estimation**

| Population | $N_{measured}$ | $N_{predicted}$ | $R^2$ | $P$ |
|---|---|---|---|---|
| FL83 | 49 | 15 | 0.52 | 2.76E-09 |
| FL81 | 31 | 27 | 0.30 | 8.13E-04 |
| AL2 | 53 | 14 | 0.77 | 5.62E-18 |
| MS6 | 55 | 17 | 0.76 | 1.88E-18 |
| AL23 | 53 | 16 | 0.45 | 1.85E-08 |
| AL22 | 52 | 14 | 0.74 | 1.16E-16 |
| AL79 | 58 | 11 | 0.74 | 2.23E-18 |
| AL21 | 45 | 18 | 0.62 | 8.67E-11 |
| MS8 | 59 | 6 | 0.71 | 3.54E-17 |
| GA1 | 46 | 15 | 0.43 | 5.39E-07 |
| ALBG | 55 | 12 | 0.58 | 1.01E-11 |
| TN3 | 65 | 9 | 0.58 | 1.04E-13 |
| TN34 | 61 | 15 | 0.51 | 5.41E-11 |
| AR2 | 57 | 11 | 0.67 | 4.75E-15 |
| KY5 | 54 | 11 | 0.68 | 7.36E-15 |
| KY1 | 40 | 11 | 0.47 | 5.60E-07 |
| MO2 | 52 | 14 | 0.71 | 2.25E-15 |
| OH1 | 60 | 8 | 0.52 | 5.62E-11 |
| KS60 | 62 | 11 | 0.61 | 5.86E-14 |
| IN5 | 55 | 10 | 0.59 | 3.52E-12 |
| OH119 | 57 | 6 | 0.63 | 8.07E-14 |
| IN4 | 53 | 17 | 0.62 | 2.18E-12 |
| IA19 | 43 | 15 | 0.57 | 5.38E-09 |

Summary statistics include the replication ($N$) of individuals with measures of reproductive output used to predict this trait for individuals without measures, the coefficient of determination ($R^2$) and the significance ($P$) of the relationship between reproductive output and stem diameter for each population, across gardens.

*Climate variables*

We calculated 14 climate variables that captured variation in temperature and precipitation at key life stages. For each common garden, daily mean, minimum and maximum temperatures (°C) were recorded during the experiment with dataloggers (temperature), and nearby weather stations (precipitation, <50km away, https://www.ncdc.noaa.gov, accessed 05/06/2024). For each population, we obtained daily temperature and precipitation data between 2018 and June 2023 from the PRISM using the *prism* R package (Edmund & Bell, 2015), based on the populations' coordinates.

We defined three time periods (hereafter "seasons", Method S3 Table D) to capture the effect of climate on key life stages in the local adaptation experiment. *Winter,* experienced by the winter cohort, was the time between the date of acclimation (or transplant for CG-Rear) and the first visit in early spring, capturing survival over winter and cueing of reproduction (i.e. vernalization). *Spring,* experienced by the summer cohort, was the time between the date of acclimation until first flowering, capturing vegetative growth and the development of the flowering stalk and buds. *Summer,* also experienced by the summer cohort, was the time between first flowering and the end of the experiment capturing reproduction.

For each season we calculated averages of daily mean, minimum and maximum temperature, and the sum of daily precipitation. We also calculated the number of days of frost and of vernalization (Method S3 Table E). These time periods were calculated for each common garden using the dates in Method S3 Table D. The seasons were also calculated for each population for each of the five years before the start of the experiment (2018 – 2022, for winter calculated from fall of each year to spring of year + 1), then averaged across years.

We also calculated five bioclimatic variables representing seasonal variation (Method S3 Table E) using the function *biovars* of the R package *dismo* (Hijmans et al., 2024) based on monthly averages of daily minimum and maximum temperatures, and monthly sum of daily precipitation. For gardens, we calculated monthly averages using a one-year daily dataset, combining temperature (from dataloggers) and precipitation (from weather station) data recorded during the season as defined in Method S3 Table D, and complemented by temperature and precipitation data obtained from weather stations to bridge gaps. For populations, we obtained daily temperatures and precipitation from PRISM to calculate monthly averages for each year between 2018 and 2022, which were then averaged across years.

**Method S3 Table D: Schedule for seasonal climate variables**

| Season | Start | End |
|---|---|---|
| *Winter 2023-2024* | *Acclimation* | *First visit in spring* |
| CG-Core | 25/10/2023 | 28/03/2024 |
| CG-Int | 08/11/2023 | 25/03/2024 |
| CG-Rear | 14/11/2023 | 26/03/2024 |
| Populations | 05/11/2023 | 26/03/2024 |
| *Spring 2023* | *Transplant* | *First flower* |
| CG-Core | 08/04/2023 | 07/06/2023 |
| CG-Int | 22/03/2023 | 19/05/2023 |
| CG-Rear | 01/03/2023 | 09/05/2023 |
| Populations | 22/03/2023 | 22/05/2023 |
| *Summer 2023* | *First flower* | *Last visit* |
| CG-Core | 07/06/2023 | 07/09/2023 |
| CG-Int | 19/05/2023 | 04/09/2023 |
| CG-Rear | 09/05/2023 | 14/08/2023 |
| Populations | 22/05/2023 | 29/08/2023 |

First flower date was the earliest date within a garden at which plants of the summer cohort opened their first flowers. The date of opening of the first flower was inferred for each plant based on the development of the flowering stalk (as in Perrier et al., 2025). Dates for populations were calculated as the average date across the common gardens.

**Method S3 Table E: Description of climatic variables**

| Variables | Description |
|---|---|
| *Seasonal climatic variables* | |
| Tmean | Seasonal average of daily mean temperatures |
| Tmin | Seasonal average of daily minimum temperatures |
| Tmax | Seasonal average of daily maximum temperatures |
| PPTsum | Seasonal sum of daily precipitation |
| DayFrost | Number of days during winter where Tmin < 0°C |
| DayVerna | Number of days during winter where Tmean < 4°C |
| *Bioclimatic variables* | |
| Bio 2 | Mean diurnal range (mean of monthly (max. temp. – min. temp.)) |
| Bio 3 | Isothermality ((Bio 2 / Bio 7) ×100) |
| Bio 4 | Temp. seasonality (standard deviation of monthly mean temp. ×100) |
| Bio 7 | Temp. annual range (max. temp. of warmest month – min. temp. of coldest month) |
| Bio 15 | Precipitation seasonality (coefficient of variation of monthly total precipitation) |

Tmean, Tmin, Tmax and PPTsum were calculated for each season (winter, spring, summer), DayFrost and DayVerna only for winter.
