## Supplementatry Tables and Figures for "The legacy of past climate warming: strong local adaptation in rear-edge populations"

**Supplementary Tables**

**Table S1: Population genetics study (Study 1) – Populations used and summary statistics**

| Pop. | Latitude [°N] | Longitude [°E] | Sequenced individuals | $\Theta_\pi$ ($\times 10^3$) | $\Theta_W$ ($\times 10^3$) | Differentiation ($F_{ST}$ ~ geography) |
|---|---|---|---|---|---|---|
| FL83 | 30.56468 | -84.95982 | 9 | 6.8089 | 6.5403 | 0.0933 |
| FL81 | 30.81165 | -85.22535 | 10 | 6.7192 | 6.6644 | 0.1704 |
| AL2 | 31.54792 | -87.51453 | 10 | 7.4527 | 7.7212 | 0.0127 |
| MS6 | 31.99803 | -89.35603 | 10 | 6.3271 | 6.3735 | 0.0994 |
| AL23 | 32.19395 | -86.78484 | 10 | 7.4147 | 7.8436 | 0.0109 |
| AL8 | 32.25610 | -86.49330 | 8 | 7.1251 | 7.0367 | 0.0670 |
| AL22 | 32.50376 | -87.50489 | 9 | 7.3975 | 7.3661 | 0.0248 |
| AL19 | 32.60699 | -88.19192 | 10 | 6.8132 | 7.0820 | 0.0794 |
| AL79 | 32.92932 | -88.20820 | 8 | 7.3443 | 7.5878 | -0.0168 |
| AL21 | 33.49109 | -86.79829 | 9 | 7.4240 | 7.2680 | 0.0211 |
| MS9 | 33.92778 | -89.01710 | 10 | 6.0470 | 5.8701 | 0.1431 |
| MS8 | 34.40427 | -88.83093 | 10 | 7.2869 | 7.3836 | -0.0100 |
| GA1 | 34.60074 | -84.69664 | 10 | 7.2319 | 7.2683 | 0.0085 |
| ALBG | 34.65346 | -86.51638 | 7 | 6.9523 | 6.6850 | 0.0664 |
| TN3 | 35.30932 | -90.06770 | 9 | 7.3577 | 7.9155 | -0.0533 |
| TN34 | 36.08222 | -86.29611 | 10 | 7.7173 | 8.7418 | -0.0719 |
| AR2 | 36.15235 | -94.30464 | 9 | 6.1749 | 6.2712 | 0.0374 |
| KY5 | 37.36005 | -84.77186 | 9 | 7.5310 | 9.2381 | -0.1045 |
| KY1 | 38.09215 | -84.98688 | 10 | 7.5346 | 8.5197 | -0.0703 |
| MO2 | 38.83025 | -92.28509 | 10 | 7.4832 | 8.7379 | -0.0990 |
| KS60 | 39.04742 | -95.68152 | 9 | 7.2858 | 7.7920 | -0.0768 |
| IN5 | 39.14583 | -86.54833 | 9 | 6.7762 | 7.0198 | 0.0031 |
| IN7 | 39.87011 | -86.16020 | 10 | 7.5731 | 8.4827 | -0.0948 |
| OH119 | 39.88500 | -83.99700 | 9 | 7.3589 | 7.7707 | -0.0536 |
| IL10 | 40.61750 | -89.01778 | 10 | 7.0531 | 7.5470 | -0.0500 |
| IN2 | 41.01731 | -85.23872 | 10 | 6.7843 | 6.9255 | -0.0167 |
| IA12 | 41.69417 | -93.67417 | 8 | 7.2769 | 7.8771 | -0.0929 |
| IA17 | 42.46194 | -90.63611 | 10 | 7.0473 | 7.6022 | -0.0531 |
| MI2 | 42.62369 | -85.43706 | 10 | 5.7276 | 5.9761 | 0.1070 |
| MN8 | 44.02861 | -92.43222 | 10 | 6.4514 | 6.8711 | -0.0055 |
| MN117 | 44.90100 | -93.19200 | 10 | 6.9598 | 7.3902 | -0.0753 |

Differentiation represents the population-average residuals of the relationship between linearized pairwise $F_{ST}$ estimates and pairwise geographic distances (Method S1).

**Table S2A: Drift load study (Study 2) – Populations used, sampling for parental and within-population cross (WPC) seed families, and population level heterosis**

| Population | Latitude [°N] | Longitude [°W] | Seed families (parental) | Seed families (WPC) | Individuals (WPC) | Drift load (average heterosis) | $F_{ST}$ (population-average) |
|---|---|---|---|---|---|---|---|
| FL83 | 30.56468 | -84.95982 | 17 | 14 | 20 | - | - |
| FL81 | 30.81165 | -85.22535 | 16 | 13 | 19 | - | - |
| **AL2** | 31.54793 | -87.51453 | 17 | 15 | 25 | -0.0590 | 0.3640 |
| **MS6** | 31.99803 | -89.35603 | 17 | 14 | 25 | 0.0842 | 0.4495 |
| **AL23** | 32.19395 | -86.78484 | 14 | 14 | 23 | 0.0350 | 0.3578 |
| AL8 | 32.25610 | -86.49330 | 18 | 15 | 20 | - | - |
| AL22 | 32.50376 | -87.50489 | 13 | 8 | 20 | - | - |
| **AL79** | 32.92932 | -88.20820 | 13 | 14 | 24 | 0.0669 | 0.3260 |
| MS9 | 33.92778 | -89.01710 | 11 | 11 | 20 | - | - |
| MS8 | 34.40427 | -88.83093 | 12 | 11 | 20 | - | - |
| **GA1** | 34.60074 | -84.69664 | 11 | 12 | 28 | 0.0389 | 0.3531 |
| **ALBG** | 34.65346 | -86.51638 | 14 | 13 | 23 | 0.2409 | 0.4037 |
| **TN3** | 35.30932 | -90.06770 | 11 | 13 | 25 | 0.1072 | 0.2854 |
| **TN34** | 36.08222 | -86.29611 | 15 | 12 | 24 | 0.0618 | 0.2659 |
| **AR2** | 36.15235 | -94.30465 | 16 | 15 | 25 | 0.1555 | 0.3927 |
| **KS60** | 39.04742 | -95.68152 | 17 | 14 | 25 | 0.1578 | 0.2888 |
| **IN5** | 39.14583 | -86.54833 | 16 | 14 | 24 | 0.1894 | 0.3447 |
| OH119 | 39.88500 | -83.99700 | 12 | 10 | 18 | - | - |
| IN4 | 40.44806 | -86.93444 | 15 | 12 | 20 | - | - |
| **PA27 *** | 41.00800 | -80.08326 | 25 | 19 | 20 | 0.2150 | 0.2821 |
| **OH64 *** | 41.11500 | -81.51810 | 25 | 18 | 19 | 0.1410 | 0.2987 |
| **MI127 *** | 41.92300 | -86.58336 | 25 | 14 | 14 | 0.3355 | 0.3333 |
| **IA10 *** | 42.07300 | -93.67250 | 25 | 21 | 21 | 0.1965 | 0.6393 |
| **MI126 *** | 42.32100 | -85.34209 | 25 | 19 | 19 | 0.1820 | 0.3699 |
| **WI128 *** | 43.15000 | -90.04498 | 25 | 19 | 20 | 0.1280 | 0.5152 |
| **MN117 *** | 44.90100 | -93.19248 | 25 | 17 | 17 | 0.2340 | 0.3089 |
| **MN118 *** | 45.02600 | -95.88795 | 25 | 20 | 21 | 0.3150 | 0.3699 |

Focal populations used to estimate drift load are indicated in bold, the other eight were used as partner populations. For eight focal populations (*), drift load was calculated from data generated in Koski *et al.* (2019), and $F_{ST}$ was calculated from data generated in Koski *et al.* (2022). Drift load was calculated for each focal population as the average heterosis (based on lifetime fitness) across its two BPC lines (Table S2B).

**Table S2B: Drift load study (Study 2) – Population pairs, sampling and heterosis estimates**

| Mother population | Father population | Seed families (BPC) | Individuals (BPC) | Heterosis |
|---|---|---|---|---|
| FL83 | **MS6** | 14 | 20 | 0.1183 |
| FL81 | **AL2** | 13 | 20 | -0.0264 |
| **AL2** | AL8 | 15 | 20 | -0.0916 |
| **MS6** | **AL23** | 14 | 20 | 0.0502 |
| **AL23** | MS9 | 14 | 20 | 0.0199 |
| **AL79** | **GA1** | 14 | 19 | 0.0121 |
| MS8 | **ALBG** | 13 | 20 | 0.3601 |
| **GA1** | AL22 | 8 | 19 | 0.0658 |
| **ALBG** | **AL79** | 13 | 20 | 0.1217 |
| **TN3** | **TN34** | 11 | 20 | 0.0136 |
| **TN34** | **AR2** | 13 | 20 | 0.1100 |
| **AR2** | **TN3** | 15 | 20 | 0.2009 |
| **KS60** | IN4 | 13 | 20 | 0.1338 |
| **IN5** | **KS60** | 14 | 20 | 0.1818 |
| OH119 | **IN5** | 10 | 18 | 0.1970 |
| **PA27 *** | **IA10 *** | 20 | 20 | 0.1230 |
| **OH64 *** | **MN118 *** | 17 | 18 | 0.3230 |
| **MI127 *** | **MN117 *** | 14 | 14 | 0.5090 |
| **IA10 *** | **MI126 *** | 20 | 21 | 0.2700 |
| **MI126 *** | **WI128 *** | 17 | 17 | 0.0940 |
| **WI128 *** | **MI127 *** | 19 | 21 | 0.1620 |
| **MN117 *** | **OH64 *** | 17 | 18 | -0.0410 |
| **MN118 *** | **PA27 *** | 20 | 22 | 0.3070 |

Populations in bold are the focal populations used to estimate heterosis, the other eight were used as partner populations. Heterosis was calculated based on lifetime fitness of within and between population crosses. For eight focal populations (*), heterosis data generated in Koski *et al.* (2019).

**Table S3: Local adaptation study (Study 3) – Populations used and replication in each cohort and common garden (CG)**

| | | | | Summer cohort (2023) | | | | | | Winter cohort (2023 – 2024) | | | | | |
|---|---|---|---|---|---|---|---|---|---|---|---|---|---|---|---|
| | | | | Families | | | Individuals | | | Families | | | Individuals | | |
| ID | Latitude [° N] | Longitude [° E] | Seed type | CG-Center | CG-Int | CG-Rear | CG-Center | CG-Int | CG-Rear | CG-Center | CG-Int | CG-Rear | CG-Center | CG-Int | CG-Rear |
| FL83 | 30.56468 | -84.95982 | Crosses | 13 | 15 | 15 | 20 | 29 | 30 | 15 | 15 | 15 | 29 | 28 | 26 |
| FL81 | 30.81165 | -85.22535 | Crosses | 14 | 15 | 15 | 20 | 30 | 29 | 15 | 15 | 15 | 29 | 30 | 30 |
| AL2 | 31.54793 | -87.51453 | Crosses | 15 | 15 | 15 | 20 | 30 | 30 | 14 | 15 | 15 | 27 | 28 | 28 |
| MS6 | 31.99803 | -89.35603 | Crosses | 13 | 15 | 15 | 20 | 30 | 30 | 15 | 15 | 15 | 28 | 30 | 29 |
| AL23 | 32.19395 | -86.78484 | Crosses | 15 | 15 | 15 | 20 | 30 | 29 | 15 | 15 | 15 | 29 | 29 | 29 |
| AL22 | 32.50376 | -87.50489 | Crosses | 13 | 15 | 15 | 20 | 30 | 29 | 15 | 15 | 15 | 29 | 29 | 29 |
| AL79 | 32.92932 | -88.20820 | Crosses | 13 | 15 | 15 | 19 | 30 | 30 | 15 | 15 | 13 | 29 | 28 | 25 |
| AL21 | 33.49109 | -86.79829 | Crosses | 13 | 15 | 15 | 20 | 30 | 29 | 15 | 15 | 15 | 29 | 28 | 28 |
| MS8 | 34.40427 | -88.83093 | Crosses | 13 | 15 | 14 | 20 | 29 | 29 | 15 | 15 | 15 | 30 | 30 | 30 |
| GA1 | 34.60074 | -84.69664 | Crosses | 14 | 15 | 15 | 19 | 30 | 30 | 15 | 14 | 14 | 28 | 26 | 25 |
| ALBG | 34.65346 | -86.51638 | Crosses | 13 | 15 | 15 | 19 | 30 | 29 | 13 | 15 | 15 | 25 | 29 | 30 |
| TN3 | 35.30932 | -90.06770 | Crosses | 13 | 15 | 15 | 20 | 30 | 29 | 15 | 15 | 15 | 29 | 29 | 30 |
| TN34 | 36.08222 | -86.29611 | Crosses | 15 | 15 | 15 | 20 | 30 | 30 | 15 | 15 | 15 | 30 | 30 | 30 |
| AR2 | 36.15235 | -94.30465 | Crosses | 13 | 15 | 15 | 20 | 29 | 30 | 15 | 15 | 15 | 29 | 30 | 30 |
| KY5 | 37.36005 | -84.77186 | Field | 14 | 15 | 15 | 20 | 29 | 30 | 15 | 15 | 15 | 29 | 28 | 30 |
| KY1 | 38.09215 | -84.98688 | Field | 11 | 15 | 15 | 19 | 29 | 30 | 13 | 14 | 12 | 26 | 26 | 22 |
| MO2 | 38.83025 | -92.28509 | Field | 12 | 14 | 14 | 18 | 27 | 29 | 14 | 14 | 13 | 27 | 26 | 26 |
| OH1 | 39.03935 | -84.32774 | Field | 12 | 15 | 15 | 20 | 29 | 27 | 15 | 15 | 15 | 30 | 30 | 30 |
| KS60 | 39.04742 | -95.68152 | Crosses | 14 | 15 | 15 | 20 | 30 | 29 | 15 | 15 | 15 | 30 | 30 | 28 |
| IN5 | 39.14583 | -86.54833 | Crosses | 13 | 15 | 15 | 19 | 30 | 29 | 15 | 15 | 14 | 29 | 30 | 26 |
| OH119 | 39.88500 | -83.99700 | Crosses | 13 | 15 | 15 | 20 | 30 | 29 | 15 | 15 | 15 | 28 | 29 | 27 |
| IN4 | 40.44806 | -86.93444 | Crosses | 13 | 15 | 15 | 19 | 30 | 29 | 15 | 15 | 15 | 29 | 29 | 29 |
| IA19 | 41.56611 | -90.47500 | Crosses | 13 | 15 | 15 | 19 | 29 | 30 | 14 | 15 | 14 | 27 | 30 | 28 |

Families and individuals depict the final number of unique seed families and individuals per population that were transplanted in each garden and tracked for fitness. Only pots with successful and timely germination were transplanted (96% of 4140 pots). In CG-Center in the summer cohort, the first 10 blocks had to be later excluded due to a mudslide. In total, 3768 seedlings were transplanted and tracked over the two cohorts.

**Table S4 Local adaptation study (Study 3) – Climate variables and the first PC axis for populations and common gardens (CG)**

| | Winter | | | | | | Spring | | | | Summer | | | | | | | | | |
|---|---|---|---|---|---|---|---|---|---|---|---|---|---|---|---|---|---|---|---|---|
| | Tmean | DayFrost | Tmax | PPTsum | DayFrost | DayVerna | Tmean | Tmin | Tmax | PPTsum | Tmean | Tmin | Tmax | PPTsum | bio2 | bio3 | bio4 | bio7 | bio15 | PC1 |
| *Populations* | | | | | | | | | | | | | | | | | | | | |
| FL83 | 13.93 | 15.00 | 20.22 | 524.69 | 15.00 | 3.00 | 21.69 | 15.06 | 28.31 | 154.39 | 27.47 | 22.04 | 32.90 | 542.59 | 12.03 | 41.56 | 628.11 | 28.94 | 28.03 | 5.83 |
| FL81 | 13.83 | 12.80 | 19.97 | 471.12 | 12.80 | 3.00 | 21.70 | 15.42 | 27.98 | 180.03 | 27.59 | 22.40 | 32.77 | 521.06 | 11.68 | 40.79 | 636.11 | 28.62 | 26.86 | 5.76 |
| AL2 | 12.05 | 36.40 | 18.84 | 484.74 | 36.40 | 9.00 | 20.01 | 12.70 | 27.33 | 177.67 | 26.63 | 20.99 | 32.28 | 458.20 | 12.99 | 42.40 | 675.34 | 30.64 | 26.79 | 4.22 |
| MS6 | 11.22 | 36.40 | 17.63 | 670.41 | 36.40 | 12.20 | 19.57 | 12.85 | 26.29 | 263.80 | 26.63 | 21.05 | 32.21 | 415.68 | 12.41 | 39.60 | 712.46 | 31.35 | 30.49 | 3.78 |
| AL23 | 11.30 | 36.20 | 17.68 | 577.98 | 36.20 | 11.60 | 19.54 | 12.72 | 26.37 | 156.19 | 26.64 | 21.12 | 32.17 | 391.93 | 12.30 | 39.59 | 708.14 | 31.07 | 14.81 | 3.54 |
| AL22 | 10.74 | 40.60 | 17.14 | 627.51 | 40.60 | 16.40 | 19.35 | 12.63 | 26.07 | 191.60 | 26.56 | 20.91 | 32.21 | 435.92 | 12.47 | 39.46 | 731.88 | 31.61 | 22.74 | 3.40 |
| AL79 | 10.30 | 39.80 | 16.48 | 743.63 | 39.80 | 17.40 | 19.18 | 12.73 | 25.63 | 230.98 | 26.57 | 21.07 | 32.07 | 410.90 | 12.04 | 37.90 | 749.24 | 31.77 | 24.23 | 3.19 |
| AL21 | 10.45 | 30.20 | 15.59 | 757.26 | 30.20 | 18.00 | 19.27 | 13.65 | 24.88 | 220.04 | 26.34 | 21.55 | 31.13 | 443.60 | 10.32 | 34.34 | 740.00 | 30.07 | 25.82 | 2.94 |
| MS8 | 9.03 | 44.20 | 14.83 | 751.16 | 44.20 | 26.60 | 18.96 | 12.93 | 25.00 | 205.60 | 26.75 | 21.39 | 32.12 | 366.89 | 11.59 | 35.04 | 817.35 | 33.08 | 30.54 | 2.21 |
| GA1 | 8.28 | 51.20 | 14.11 | 646.62 | 51.20 | 28.20 | 17.52 | 10.89 | 24.16 | 146.76 | 25.24 | 19.61 | 30.86 | 333.53 | 11.99 | 36.93 | 773.44 | 32.48 | 24.77 | 0.95 |
| ALBG | 7.98 | 53.40 | 13.52 | 803.97 | 53.40 | 34.00 | 17.36 | 11.28 | 23.43 | 205.16 | 25.05 | 19.78 | 30.33 | 375.49 | 11.37 | 35.25 | 787.57 | 32.26 | 28.87 | 0.86 |
| TN3 | 7.67 | 44.80 | 12.73 | 675.94 | 44.80 | 34.40 | 18.46 | 13.02 | 23.91 | 193.33 | 26.59 | 21.58 | 31.59 | 274.90 | 10.60 | 31.77 | 871.38 | 33.37 | 34.51 | 0.84 |
| TN34 | 6.84 | 69.40 | 13.20 | 706.96 | 69.40 | 41.80 | 16.24 | 9.11 | 23.38 | 189.93 | 25.01 | 18.70 | 31.32 | 400.29 | 13.18 | 37.70 | 829.34 | 34.97 | 26.60 | 0.26 |
| AR2 | 5.37 | 75.20 | 11.21 | 408.07 | 75.20 | 55.20 | 16.01 | 10.29 | 21.73 | 277.09 | 24.88 | 19.47 | 30.29 | 304.75 | 11.55 | 32.66 | 896.39 | 35.37 | 39.55 | -1.45 |
| KY5 | 4.65 | 76.40 | 10.05 | 639.24 | 76.40 | 62.20 | 14.93 | 8.74 | 21.12 | 186.46 | 23.50 | 17.89 | 29.11 | 424.26 | 11.56 | 33.74 | 858.56 | 34.26 | 20.69 | -1.84 |
| KY1 | 3.97 | 80.20 | 9.12 | 518.98 | 80.20 | 68.20 | 14.69 | 8.90 | 20.48 | 157.17 | 23.58 | 18.15 | 29.01 | 387.16 | 11.06 | 31.68 | 895.37 | 34.90 | 20.17 | -2.66 |
| MO2 | 2.64 | 87.60 | 8.03 | 331.60 | 87.60 | 73.60 | 15.09 | 9.32 | 20.85 | 178.42 | 24.70 | 19.08 | 30.33 | 315.03 | 11.44 | 30.22 | 1006.40 | 37.85 | 35.34 | -3.28 |
| OH1 | 3.18 | 86.00 | 8.13 | 441.33 | 86.00 | 74.80 | 14.25 | 8.44 | 20.06 | 179.55 | 23.43 | 18.02 | 28.84 | 384.76 | 10.78 | 30.80 | 922.27 | 35.00 | 22.41 | -3.29 |
| KS60 | 2.07 | 103.80 | 8.11 | 194.38 | 103.80 | 80.00 | 14.91 | 8.93 | 20.89 | 181.55 | 25.09 | 19.55 | 30.62 | 336.89 | 12.02 | 30.58 | 1042.56 | 39.32 | 61.91 | -3.79 |
| IN5 | 2.49 | 93.00 | 7.45 | 494.78 | 93.00 | 81.00 | 13.94 | 7.99 | 19.89 | 158.25 | 23.34 | 17.76 | 28.92 | 380.28 | 11.05 | 30.59 | 956.30 | 36.13 | 21.21 | -3.76 |
| OH119 | 1.74 | 96.60 | 6.57 | 402.92 | 96.60 | 88.20 | 13.30 | 7.63 | 18.96 | 160.35 | 22.94 | 17.49 | 28.40 | 323.92 | 10.73 | 29.88 | 965.83 | 35.90 | 22.42 | -4.67 |
| IN4 | 0.45 | 102.20 | 5.00 | 315.31 | 102.20 | 96.60 | 12.86 | 7.17 | 18.55 | 150.08 | 22.73 | 17.07 | 28.38 | 261.93 | 10.73 | 28.96 | 1019.50 | 37.06 | 27.80 | -5.78 |
| IA19 | -1.56 | 113.60 | 2.99 | 245.64 | 113.60 | 106.40 | 12.02 | 6.58 | 17.46 | 163.73 | 22.87 | 17.68 | 28.05 | 271.78 | 10.19 | 25.56 | 1113.19 | 39.86 | 40.63 | -7.27 |
| *Common gardens* | | | | | | | | | | | | | | | | | | | | |
| CG-Rear | 13.63 | 4.00 | 20.51 | 698.13 | 4.00 | 2.00 | 19.23 | 13.91 | 25.32 | 180.33 | 25.68 | 21.87 | 30.61 | 464.22 | 11.04 | 38.38 | 588.13 | 28.76 | 45.95 | 4.35 |
| CG-Int | 8.50 | 79.00 | 16.07 | 707.66 | 79.00 | 27.00 | 16.68 | 10.77 | 24.12 | 285.21 | 22.94 | 18.76 | 28.67 | 493.39 | 12.71 | 39.25 | 711.61 | 32.37 | 57.53 | 0.72 |
| CG-Center | 6.46 | 69.00 | 14.15 | 514.40 | 69.00 | 53.00 | 16.42 | 10.33 | 22.94 | 122.99 | 22.14 | 18.32 | 26.77 | 392.85 | 11.95 | 37.67 | 716.13 | 31.72 | 47.85 | -1.16 |

Variable description and calculation detailed in Method S3

**Table S5 Local adaptation study (Study 3) – Summary of the principal component analysis (PCA) on seasonal and bioclimatic variables**

| | | PC1 | PC2 | PC3 | PC4 |
|---|---|---|---|---|---|
| Variance explained (%) | | 76.65 | 8.81 | 5.86 | 4.59 |
| Cumulative variance (%) | | 76.65 | 85.46 | 91.32 | 95.9 |
| *Loadings* | | | | | |
| Winter | Tmean | **0.26** | -0.02 | 0.00 | 0.02 |
| | Tmin | **0.26** | -0.05 | -0.06 | 0.09 |
| | Tmax | **0.26** | 0.00 | 0.05 | -0.04 |
| | PPTsum | 0.16 | **-0.22** | **-0.51** | **-0.31** |
| | DayFrost | **-0.26** | 0.04 | 0.10 | -0.13 |
| | DayVerna | **-0.26** | 0.00 | 0.04 | 0.03 |
| Spring | Tmean | **0.26** | 0.08 | 0.00 | 0.12 |
| | Tmin | 0.25 | 0.11 | -0.10 | 0.24 |
| | Tmax | **0.26** | 0.05 | 0.08 | 0.02 |
| | PPTsum | 0.08 | **0.40** | **-0.54** | **-0.49** |
| Summer | Tmean | 0.25 | **0.23** | 0.02 | 0.13 |
| | Tmin | 0.24 | **0.24** | -0.08 | **0.25** |
| | Tmax | 0.25 | 0.21 | 0.12 | 0.01 |
| | PPTsum | 0.20 | -0.21 | **0.24** | 0.02 |
| Bioclim. | Bio 2 (Mean diurnal range) | 0.15 | 0.08 | **0.50** | **-0.64** |
| | Bio 3 (Isothermality) | 0.25 | -0.08 | **0.23** | -0.22 |
| | Bio 4 (Temperature seasonality) | -0.25 | 0.16 | -0.01 | 0.05 |
| | Bio 7 (Temperature annual range) | -0.25 | 0.20 | 0.11 | -0.11 |
| | Bio 15 (Precipitation seasonality | 0.08 | **0.70** | 0.14 | 0.10 |

Data only shown for principal components (PC) 1 to 4, as the cumulative contribution of the remaining PCs explained less than 5% of the variance. Loadings highlighted in bold indicate the main contributing variables to a given PC axis (upper quartile based on absolute loading for each PC).

**Table S6: Model comparisons between testing linear and quadratic effects of latitude**

| Dependent variable | *N* | *AICc* linear | *AICc* square | $R^2$ linear | $R^2$ square |
|---|---|---|---|---|---|
| *Study 1 - Population genetics* | | | | | |
| $\theta_W$ | 31 | -490.82 | **-494.00** | 0.00 | 0.14 |
| $\theta_\pi$ | 31 | -521.84 | -522.88 | -0.02 | **0.06** |
| Differentiation | 31 | -78.82 | **-83.95** | 0.30 | 0.44 |
| *Study 2 - Drift load* | | | | | |
| Heterosis | 19 | **-43.33** | -40.51 | 0.54 | 0.53 |

The best model (bold) was identified by the lowest AICc value. For models with similar AICc (|ΔAICc|<2), the best model was the one with highest $R^2$ (bold).

**Table S7A: Local adaptation study (Study 3) – Comparison of relative lifetime fitness between common gardens (CG)**

| | $\beta$ Common gardens | |
|---|---|---|
| CG-Center vs CG-Int | -0.04 | |
| CG-Center vs CG-Rear | **0.24** | ** |
| CG-Int vs CG-Rear | **0.28** | *** |

Values depict the estimated pairwise differences ($\beta$) between common gardens obtained from Tukey's tests. Estimates with *P*-values < 0.05 in bold; ** $P$<0.01, *** $P$<0.001.

**Table S7B: Local adaptation study (Study 3) – Estimates of the effect of ΔPC1 on relative lifetime fitness**

| | $\beta$ ΔPC1 | $\beta$ $\Delta PC1^2$ |
|---|---|---|
| CG-Center | 0.024 | -0.003 |
| CG-Int | 0.003 | -0.003 |
| CG-Rear | -0.103 | 0.003 |

Values depict the estimated slope coefficients ($\beta$) of the linear and quadratic effect of ΔPC1 in each common garden (CG).

**Table S7C: Local adaptation study (Study 3) – Comparison of the effect of ΔPC1 on relative lifetime fitness between common gardens (CG)**

| | $\beta$ Common gardens * ΔPC1 | |
|---|---|---|
| CG-Center vs CG-Int | 0.02 | |
| CG-Center vs CG-Rear | **0.11** | ** |
| CG-Int vs CG-Rear | **0.09** | *** |

Values depict the estimated pairwise contrast ($\beta$) of the effect of ΔPC1between common gardens (based on the garden - ΔPC1 interaction) obtained from Tukey's tests. Estimates with *P*-values < 0.05 in bold; ** $P$<0.01, *** $P$<0.001.

**Table S8A: Local adaptation study (Study 3) – Test of variation in lifetime fitness across common gardens and population latitude.**

| Fixed effects | $X^2$ | |
|---|---|---|
| Common garden (CG) | **87.92** | *** |
| Latitude | **9.36** | ** |
| CG * latitude | **136.88** | *** |

The dependent variable was population-average lifetime fitness, assumed to follow Gaussian distributions. All fixed effects were tested in one model, optimized with the bobyqa optimizer to improve convergence. Test statistics include the chi-squared value of the model, bold if P-values < 0.05; ** $P$<0.01, *** $P$<0.001. Marginal $R^2_m$ = 0.70 (fixed effect only); $R^2_c$ = 0.83 (fixed and random effects). Results for random effects are not shown.

**Table S8B: Local adaptation study (Study 3) – Comparison of lifetime fitness between common gardens (CG)**

| | $\beta$ Common gardens | |
|---|---|---|
| CG-Center vs CG-Int | -0.11 | |
| CG-Center vs CG-Rear | **0.58** | *** |
| CG-Int vs CG-Rear | **0.69** | *** |

Values depict the estimated pairwise differences ($\beta$) between common gardens obtained from Tukey's tests. Estimates with $P$-values < 0.05 in bold; *** $P$<0.001.

**Table S8C: Local adaptation study (Study 3) – Estimates of the effect of latitude on lifetime fitness**

| | $\beta$ Latitude | $\beta$ Latitude$^2$ |
|---|---|---|
| CG-Center | 0.419 | -0.005 |
| CG-Int | 0.364 | -0.005 |
| CG-Rear | -0.780 | 0.009 |

Values depict the estimated slope coefficients ($\beta$) of the linear and quadratic effect of latitude in each common garden (CG).

**Table S8D: Local adaptation study (Study 3) – Comparison of the effect of ΔPC1 on relative lifetime fitness between common gardens (CG)**

| | $\beta$ Common gardens * Latitude | |
|---|---|---|
| CG-Center vs CG-Int | **0.06** | ** |
| CG-Center vs CG-Rear | **0.21** | *** |
| CG-Int vs CG-Rear | **0.15** | *** |

Values depict the estimated pairwise contrast ($\beta$) of the effect of latitude between common gardens (based on the garden – latitude interaction) obtained from Tukey's tests. Estimates with $P$-values < 0.05 in bold; ** $P$<0.01, *** $P$<0.001.

### Supplementary Figures

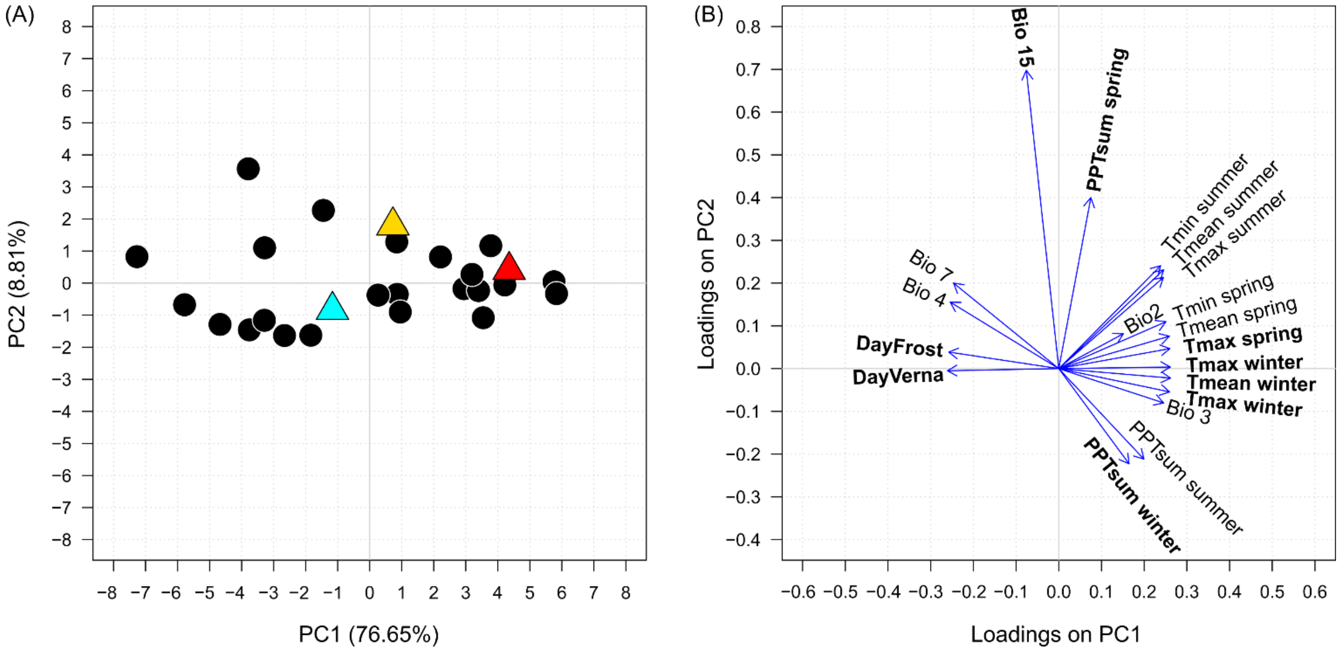

**Figure S1. Local adaptation study (Study 3) – Estimation of climate differences between gardens and populations**
**(A)** Environmental variation across the 23 populations (dots) included in the experiment is represented by their relative position along the first two axes of a principal component analysis, based on 19 climatic variables estimated for each populations' location of origin (Table S3, Method S3). Common gardens (triangles; cyan, CG-Center; yellow, CG-Int; red, CG-Rear) were projected onto the populations' PC space based on the same climate variables. **(B)** The contribution of climate variables to the first two principal components are indicated by arrows, with the main contributing variables labeled in bold (Table S5).

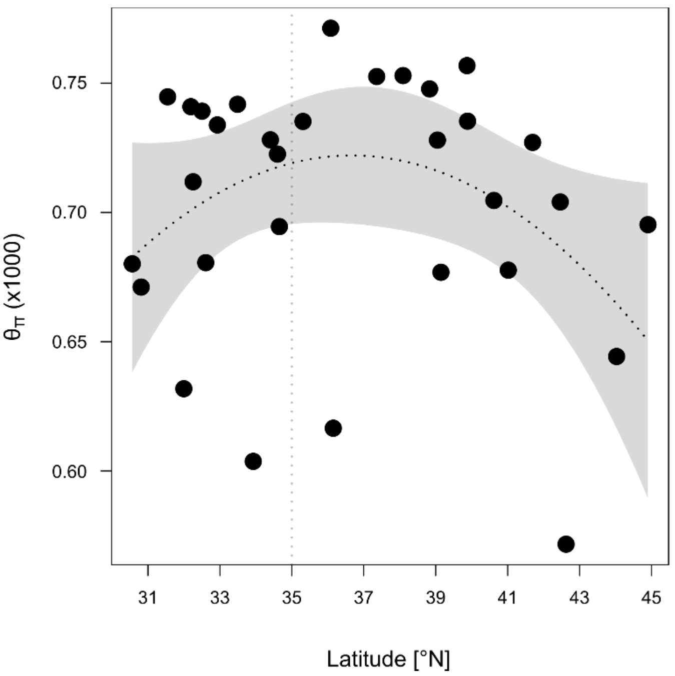

**Figure S2**: **Population genetic study (Study 1) – Variation in genetic diversity across latitudes**
Genetic diversity within population based on Tajimas' $\theta_{\pi}$ estimated for each population (dots). Vertical dotted line represents the separation between the rear edge (<35°N) and the expanded range. Dashed black lines represent the model-predicted relationship between each estimate and population latitude, with the 95% confidence interval indicated as shading. Test statistics are reported in Table 1.

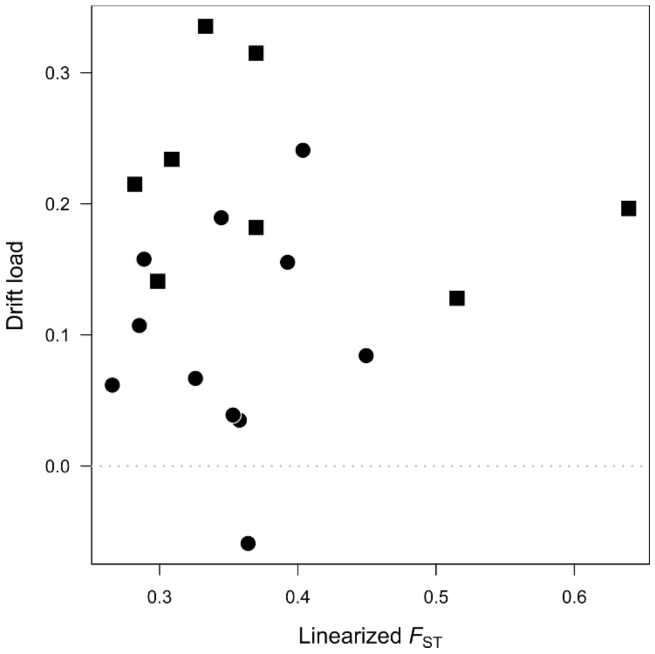

**Figure S3: Drift load study (Study 2) – Variation in the expression of drift load with population differentiation ($F_{ST}$)**

Drift load of each focal population (dots) and additional populations (squares) plotted again linearized $F_{ST}$. The lack of significant relationship indicates that drift load is not associated with genetic differentiation among populations.

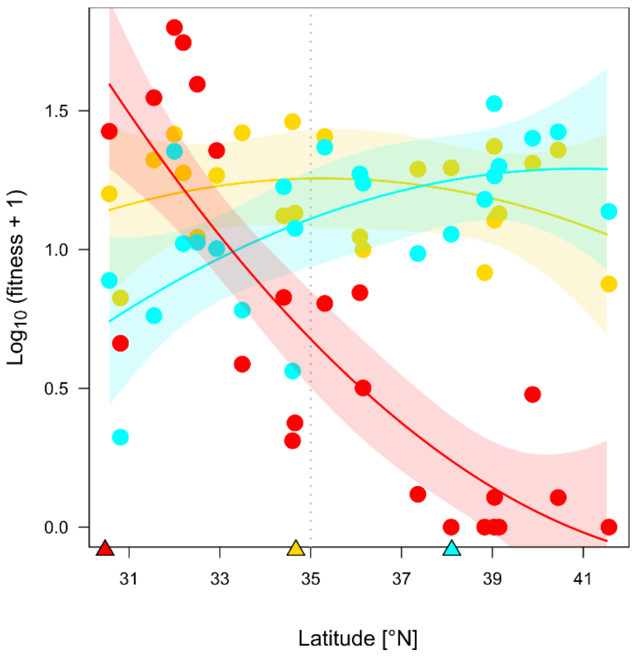

**Fig. S4 Local adaptation study (Study 3) – Variation in lifetime fitness across latitude and common gardens (CG)**

Average lifetime fitness of populations (dots) raised in each common garden (cyan, CG-Center; yellow, CG-Int; red, CG-Rear). Population averages were calculated from log10 transformed family averages (after adding +1 so that all values are positive). Vertical dotted line represents the separation between the rear edge (<35°N) and the expanded range. Solid lines represent the relationship between lifetime fitness and the populations' latitude, estimated for each garden, with the 95% CI indicated in shading. Colored triangles indicate the latitude of each garden. Test statistics are reported in Table S8.
